# Rheb-mTOR Activation Rescues Amyloid Beta-Induced Cognitive Impairment and Memory Function by Restoring miR-146 Activity in Glial Cells

**DOI:** 10.1101/2021.02.11.430878

**Authors:** Dipayan De, Ishita Mukherjee, Subhalakshmi Guha, Ramesh Kumar Paidi, Saikat Chakrabarti, Subhas C. Biswas, Suvendra N. Bhattacharyya

## Abstract

Deposition of amyloid beta plaques in adult rat or human brain is associated with increased production of proinflammatory cytokines by associated glial cells that are responsible for degeneration of the diseased tissue. The expression of these cytokines is usually under check and is controlled at post-transcriptional level via several microRNAs. Computational analysis of gene expression profiles of cortical regions of Alzheimer’s disease patients brain suggests ineffective target cytokine mRNA suppression by existing microRNPs in diseased brain. Exploring the mechanism of amyloid beta induced cytokine expression, we have identified how the inactivation of the repressive miR-146 microRNPs causes increased production of cytokines in amyloid beta exposed glial cells. In exploration of the cause of miRNP inactivation, we have noted amyloid beta oligomer induced sequestration of mTORC1 complex to early endosomes that results in decreased Ago2 phosphorylation, limited Ago2-miRNA uncoupling and retarded Ago2-cytokine mRNA interaction in rat astrocyte cells. Interestingly, constitutive activation of mTORC1 by Rheb activator restricts proinflammatory cytokine production by reactivating miR-146 microRNPs in amyloid beta exposed glial cells to rescue the disease phenotype in the *in vivo* rat model of Alzheimer’s disease.

## Introduction

MicroRNAs (miRNA) are 20-22 nucleotides long non-coding RNAs that can fine-tune the expression of their target mRNAs post transcriptionally in metazoan cells by imperfect base pairing^1, 2^. Like in other tissues, miRNAs are important gene regulators in mammalian brain, where hundreds of miRNAs are known to control thousands of transcripts encoding important proteins that affect several physiological processes and events in mammalian brain^3^. Differentiation of embryonic stem cells to neuronal cell is also controlled by specific miRNAs like miR-9^4^ while miR-128 regulates the growth of the dendrites^5^. There are also evidences of miRNA playing a significant role in neurodegenerative disease (NDDS) like in Alzheimer’s Disease (AD), Parkinson’s Disease (PD), Huntington’s Disease (HD) or in Amyotrophic Lateral Sclerosis(ALS)^6^. There have been reports which show the effect of conditional knockout of Dicer, an essential enzyme for miRNA biogenesis, on neuronal death^7^. Additionally, differential miRNA expression profiles from postmortem brains of patient died of AD have shown the possible importance of miRNA expression in pathophysiology of this disease^8, 9^.

Astrocytes are the most abundant cells in mammalian brain. The increase of number and complexity of this glial cell population during brain development support their possible role in regulating complex traits such as human cognition and behavior^10^. Astrocytes provide support to the neuronal function by regulating ion homeostasis^11^, or by controlling CNS blood flow^12^ and they also play crucial role in synaptic transmission by forming tripartite synapse^13, 14^. Apart from their supportive role, astrocytes are also immunomodulatory cells known to secrete various cytokine and chemokines which play an important role in pathophysiology of different diseases including AD^15^. In AD increased production of inflammatory cytokines may leads to death of the neighboring neurons and considered as the major path that amyloid beta (Aβ) oligomers activates in glial cell population leading to neuronal death^16^. However, the exact mechanism of the excess cytokine production in glial cells by Aβ exposure is not clear. miRNAs are important regulators of immune response^17^ and they can control hyper responsiveness of immune cells by reversibly regulating miRNA activity in activated macrophage cells^18, 19^. So expression and activity of mRNAs in astroglia, the major immune cells in brain context, likely to have a role to play in controlling the neuroinflammation during neurodegenerative diseases.

Argonaute (Ago) proteins play the crucial role in post transcriptional gene silencing in eukaryotic cells. According to published reports, Ago proteins are responsible for the regulation of expression of majority of protein coding genes primarily by associating with specific miRNAs having complementarities with those protein coding genes^20, 21^. In mammals, there are four Ago proteins (Ago1-4) and Ago2 is the major and widely expressed Ago in different tissues including brain^22^. Upon its association with miRNAs, it can cleave the target mRNAs having perfect complementarities with bound miRNAs^23^. Post translation modification of Ago protein can also determine its function. Increased Ago stability and catalytic activity are associated with modifications like hydroxylation and methylation of Ago^24, 25^. Among all the modifications, phosphorylation at different positions may play a significant role in miRNA binding of Ago2 and its sub cellular localization^26–28^.

Mammalian target of Rapamycin (mTOR), a master regulator of cell growth comprises of mTORC1 and mTORC2 which are different forms of structural and functional complexes involving mTOR protein. Raptor, mLST8 along with mTOR are the major subunits of mTORC1 which regulates cell growth, protein homeostasis and synthesis, along with autophagy^29^. mTORC2 constitutes mTOR, Rictor, SIN1 and mLST8. The major function of this complex includes the lipogenetic regulation and that of cytoskeletal structure^30^. mTOR pathway specializes in sensing multiple external stimuli such as growth factors, oxygen level and is known to respond by sensing amino acid and glucose concentration as well. Important neuronal physiological phenomenon like synaptic plasticity, memory and learning are often maintained by mTOR^31^. Faulty mTORC1 signaling has been linked with AD previously and TREM2 deficient microglial cells were shown to have decrease in mTORC1 activity that have contributed to high autophagic flux leading to weakened microglial fitness^32^. Studies have shown that in spite of known upstream and downstream regulatory factors, the major key to the mTOR associated signaling lies in the compartmentalization of the mTORC1 to specific cellular sites^33, 34^. How does disease conditions influence mTOR localization or does mTOR localization play a role in pathophysiology of a disease are the unresolved questions in the field.

In this study, we report how the exposure of Amyloid-β oligomers (Aβ_1-42_) in rat astrocyte cells decreases the cellular mTORC1 activity by enhancing the sequestration of mTORC1 complexes to early endosome. Aβ_1-42_ also restricts translocation of mTORC1 to the lysosome which is found to be important for mTORC1 activation. mTOR inactivation reduces miRNA activity due to a decrease in Ago2 phosphorylation. Interestingly, Reactivation of mTORC1 through the expression of the constitutive activator Rheb, reverses both Aβ_1-42_ mediated pro-inflammatory cytokine production and restores behavioral abnormalities in animals along with a rescue of miRNA function suggesting a pivotal role of miRNA activity modulation by mTORC1 in AD.

## Results

### Exposure to Aβ_1-42_ oligomer causes miRNA upregulation in human and adult rat brain

miRNAs are important regulators of majority of metazoan genes that are supposed to be upregulated and expressed profoundly upon inactivation or downregulation of cognate miRNAs. To obtain an insight of how the miRNA and mRNA profile gets altered in the disease process, we analyzed the “regulator(s) to target(s)” (miRNA:mRNA) expression profiles in different brain regions of Alzheimer’s disease (AD) patients. The relationship between the differentially expressed miRNAs and their corresponding target genes were studied. A set of 30 miRNAs were found to be up-regulated whereas 23 miRNA were found to be down-regulated in the pre-frontal cortex of Alzheimer’s patients (Figure 1A and Table S1) and these miRNAs are likely to be related to Alzheimer’s disease etiology or progression. clustering has been done for analyzed miRNAs. Analysis of miRNA expression profiles in pre-frontal cortex of Alzheimer’s disease patients was performed considering the samples provided in the GEO dataset (GSE48552)^35^. Based on clinical presentation and stage of the disease, the samples had been grouped into early stage and late stage Alzheimer’s. Differentially expressed miRNA between these stages were determined utilising DESeq^36^ package in R. Normalized counts of the differentially expressed miRNA identified considering these samples have been depicted in the heatmap (Figure 1A). Further, considering mRNA expression profiling data 2092, 2166 and 292 mRNA were found to be up-regulated while 1253, 699 and 676 genes were found to be down-regulated within the superior frontal gyrus, frontal cortex and dorso-lateral prefrontal cortex respectively. Moreover, 271 genes (mRNA) were found to be commonly up-regulated among two or more of the different brain cortical regions considered herein for the analysis. The corresponding fraction of up-regulated target mRNAs of each of the differentially expressed miRNA was determined in different cortical regions (frontal cortex, superior frontal gyrus and dorsolateral pre frontal cortex) of the human brain. Herein, we observed that in comparison to the down-regulated miRNAs, nearly 79% of the up-regulated miRNAs had a substantial fraction (0.6-0.8) of their target mRNAs also as up-regulated in the frontal cortex (Figure 1C, D). Further, the observation that in comparison to the down-regulated miRNAs, a higher percentage of the up-regulated miRNAs have a substantial fraction of their target mRNAs as up-regulated is consistent within the other cortical regions considered in this study. Herein, in comparison to the down-regulated miRNAs, 88% and 48% of the up-regulated miRNAs had between 0.4-0.6 or 0.2-0.4 fraction of their target mRNA as up-regulated within the superior frontal gyrus and dorsolateral pre-frontal cortex, respectively (Figure 1C, D). In particular, the up-regulated miRNA (~65%) and down-regulated miRNA (~67%) have similar fraction (between 0.4-0.6 or 0.2-0.4) of their corresponding target mRNAs as up-regulated in the cancer case studies considered here (Figure 1E and in additional information provided as Appendix).

**Figure 1.**
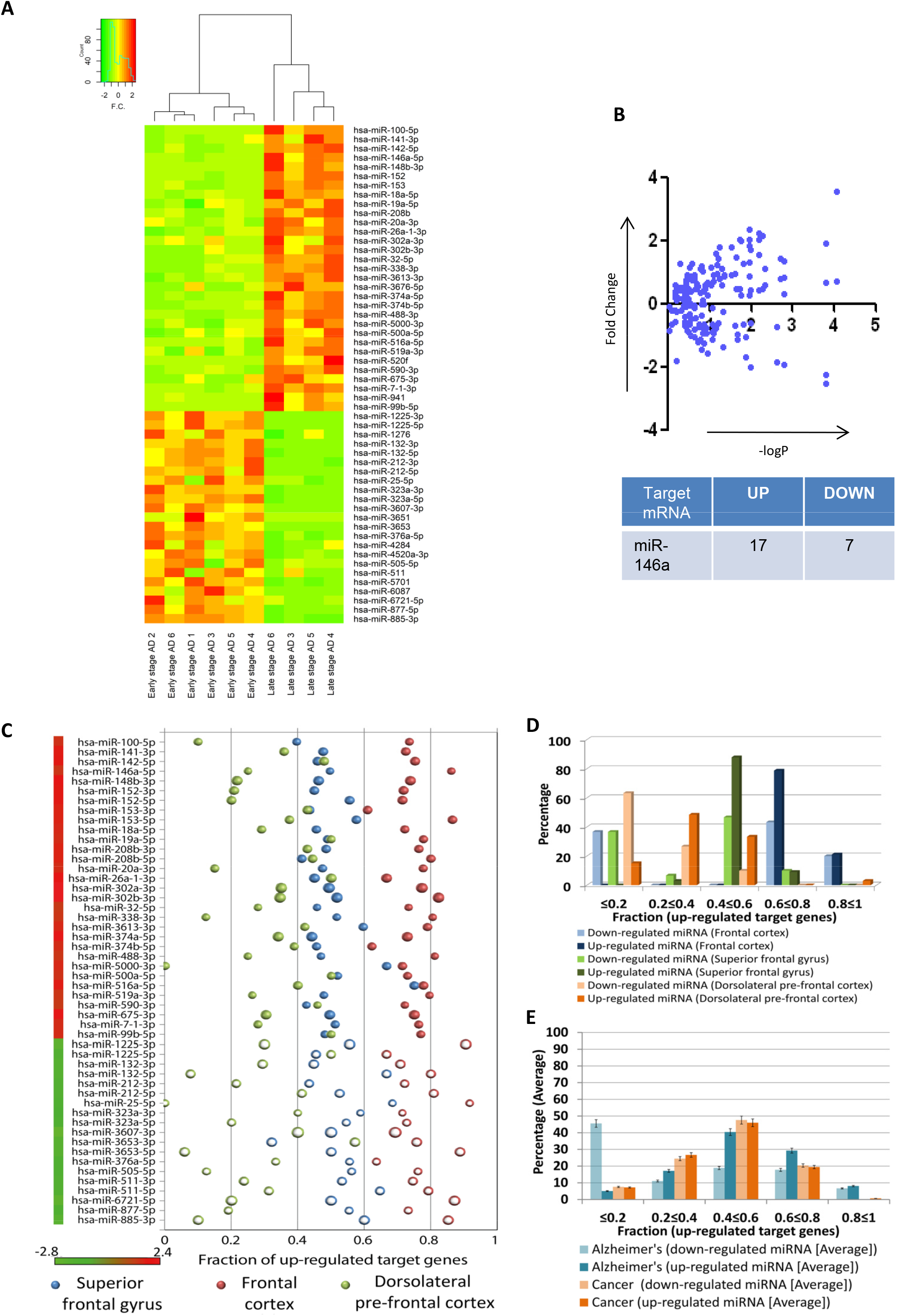
Figure 1 Possible miRNA to mRNA expression relationships in Alzheimer’s disease and cancer. **A** Heatmap of differentially expressed miRNAs in pre-frontal cortex of Alzheimer’s disease patients. The significance threshold for selection of differentially expressed miRNA was considered as fold change is +/-1.25 and of p-value <= 0.05. **B** Graphs showing expression level of miR-146a targets in pre-frontal cortex of Alzheimer’s disease patients. Y axis represents fold change of individual mRNAs and X axis represents −log_10_P value. The table represents number of miR-146a targets having at least 1.5 fold changes. **C** The probable relationship between differentially expressed miRNA and mRNA in Alzheimer’s disease considering the possible fraction of up-regulated target mRNA of each of the miRNA as studied in different cortical regions (frontal cortex, superior frontal gyrus and dorsolateral pre-frontal cortex) has been depicted here. **D** Percentages of up-regulated or down-regulated miRNA that possibly have a fraction of their target genes as up-regulated within the frontal cortex, superior frontal gyrus and dorsolateral pre-frontal cortex is shown here. **E** Comparison among percentage of differentially expressed miRNA that have different subsets or fractions of their target mRNA as up-regulated in degenerative tissues (AD) as opposed to highly proliferative tissues (Cancer) has been performed.

Importantly, we have considered multiple datasets to identify a common trend in the de-regulated miRNA-mRNA interaction network by considering target mRNA that are commonly differentially expressed in two or more of the different brain cortical regions. However, during the analysis we have considered all the target mRNA that were significantly differentially expressed in each dataset to study the “regulator(s) to target(s)” (miRNA:mRNA) expression profile patterns in Alzheimer’s disease (AD) patients and compared the same. miR-146a-5p has consistently been found to exhibit this phenomenon across all the conditions analyzed herein. Further, miR-146a which is known to regulate expression of number of cytokines^37^, was found to be upregulated. Expression of all the probable miR-146a targets was also analyzed. Interestingly, majority of the miR-146a targets were found to be upregulated in AD brain indicated a possible loss of function of miR-146a on its target in AD brain (Figure 1B and Table S2).

In the Alzheimer’s disease datasets considered here for analysis, a substantial percentage of up-regulated miRNA have a large fraction of their corresponding target genes as up-regulated. This is in contrary to the general regulatory relationship between miRNA and their target genes wherein the target genes of up-regulated miRNA are down-regulated as expected^2, 38^. However, this inverse relationship between the target genes and up-regulated miRNA has also been noted in other cell types other than degenerative brain. In cancer cells, the trend of target gene upregulation in the backdrop of up-regulated cognate miRNA expression has also been noted (see Appendix and Figure S1). However, the upregulation of target gene with increase in cognate miRNAs has been more prominent in AD rather than in cancer (Figure 1D-1E). Thus, we observed trends that in comparison to the down-regulated miRNA, target mRNA of up-regulated miRNA are mainly up-regulated and this phenomenon is more prominent in Alzheimer’s disease condition than in cancer. Therefore in AD brain there has been a possibility of existence of a mechanism for gross inactivation of miRNPs that may let them ineffective to suppress their targets.

We wanted to explore the effect of Aβ_1-42_ oligomer, the suspected causative link between amyloid plaque and the disease phenotype in Alzheimer’s patient, on cellular miRNA levels in rat brain. The pre-formed Aβ_1-42_ oligomer was stereotactically injected in adult rat brain hippocampal region and animal were kept for 21 days which is known to cause deposition of Aβ plaques in and around the injection area (Figure 2A-B)^39^. A deposition of Aβ plaques also resulted in GFAP activation which is a marker for astrocyte proliferation as well as immune activation in rat brain (Figure 2B)^40^. Now there are several miRNAs present in rat brain which can regulate the expression profile of these cytokine mRNAs either directly or indirectly. For example, IL-6 is a direct target of Let-7a^41^ and miR-146a fine tunes cytokine production by repressing NF-κB pathway^42^. So we wanted to check what is expression pattern of those miRNAs which directly and indirectly regulates expression of pro-inflammatory cytokines in both rat brain as well as in primary rat astrocytes. mRNA levels of different proinflammatory cytokines like IL-1β, IL-6 were increased several fold when oligomeric Aβ_1-42_ was stereotactically infused in the hippocampus of adult rat brain supporting already reported Aβ_1-42_ induced immunoactivation reaction in affected brain (Figure 2C). Interestingly, as observed in human patient samples, expression profile of several miRNAs including the immunogenic miRNAs miR-21, miR-146a, and miR-155 also showed an upregulation in adult rat brain injected with Aβ_1-42_ (Figure 2D). To identify the contribution of astrocyte cells in cytokine production in rat brain upon activation with Aβ_1-42_ oligomer, we treated the primary rat astrocyte cells with Aβ_1-42_ oligomer *ex vivo*. The activation of the cells was scored by excess production of GFAP in rat primary astrocyte on Aβ_1-42_ oligomer exposure (Figure 2E-F). It was found that Aβ can activate astrocyte cells where mRNA level of different cytokines were found to be increased several fold (Figure 2E-F). We noted a global increase in miRNA level in Aβ_1-42_ treated astrocyte cells where 65 to 70 % of the miRNAs were found to be upregulated including miR-146a, miR-155, miR-21 and Let-7a. These miRNAs are direct and indirect regulators of IL-1β. IL-6 also noted to be increased in treated astrocyte (Figure 2G). Interestingly we found increase in several other miRNAs such as miR-16, miR-29a, miR-145 in both brain tissue as well as in primary glial cells upon amyloid exposure (Figure 2D, 2G). These miRNAs are not known to affect cytokine expression in mammalian cells, indicating the possibility that the phenomenon we are observing could have broader implications and affecting several different types of miRNAs.

**Figure 2.**
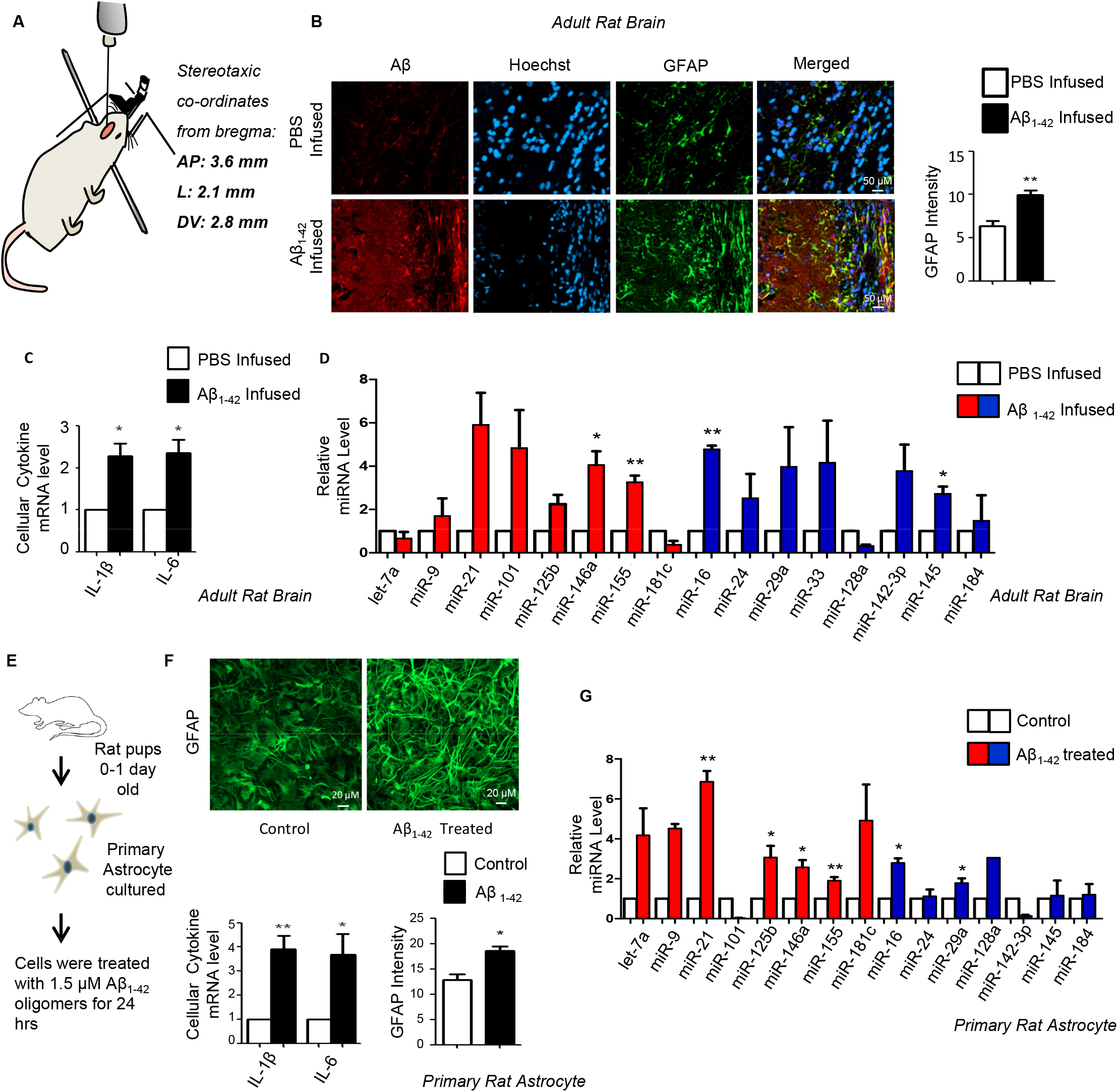
Aβ_1-42_ induces inflammatory response and increases miRNA level in astrocyte cells. **A** Preformed Aβ_1-42_ oligomers in PBS was infused in the hippocampus at stereotaxic co-ordinates from bregma: AP:2.6, L:2.1, DV:2.8 mm in adult rat brain. PBS without Aβ_1-42_ was used for control injection. Animals were kept 21days for plaque formation. **B** Cryosections of Aβ_1-42_ infused and PBS infused control rat brain hippocampus were subjected to immunohistochemistry (IHC) analysis. Astrocytes were immunostained for GFAP (green) whereas Aβ was visualized in red and nucleus was stained with Hoechst (blue). Relative intensity of the GFAP signals were measured by ImageJ software and quantification has been shown in the right panel. **C** qPCR based quantification of total mRNA levels of IL-1β and IL-6 in PBS and Aβ_1-42_ infused adult rat hippocampus. The data was normalized against GAPDH mRNA level. **D** Cellular miRNA levels in PBS and Aβ_1-42_ infused adult rat brain hippocampus. Taqman based miRNA level quantification was done. Values were normalized with U6 snRNA level. Red represents the miRNAs those are known to regulate cytokine mRNAs and blue represents miRNAs which are not known to regulate cytokine mRNAs. **E-F** Experimental strategy of isolation of rat primary astrocyte from 0-1 day old rat pup brain followed by treatment of the primary cells with 1.5 μM Aβ_1-42_ oligomers *ex vivo* (E). Microscopic images showing effect on Aβ_1-42_ on primary cortical astrocytes stained with GFAP (Green, F *upper panel*).The intensity of the GFAP in both conditions were calculated by ImageJ software and plotted (F, *bottom right panel*). Graphs depicting levels of cellular IL-1β and IL-6 mRNA level in Aβ_1-42_ treated primary cortical astrocytes. The data was normalized against GAPDH mRNA level (F, *lower left panel*). **G** Cellular miRNA levels in Aβ_1-42_ treated primary cortical astrocytes. Taqman based miRNA level quantification were done. Values were normalized with U6 snRNA. Red represents the miRNAs those are known to regulate cytokine mRNAs and blue represents miRNAs which are not known to regulate cytokine mRNAs. For statistical significance, minimum three independent experiments were considered in each case unless otherwise mentioned and error bars are represented as mean ± S.E.M. P-values were calculated by utilizing Student’s t-test. ns: non-significant, *P < 0.05, **P < 0.01, ***P < 0.0001.

### mTOR mediated Ago2 phosphorylation is required for miRNA-mediated repression of targets

To uncover the mechanism of non-functional miRNP accumulation in Aβ_1-42_ treated cells, we used C6 glioblastoma cells to treat with Aβ_1-42_ oligomer before the biochemical analysis was done for the treated and control cells. To confirm the accumulation of non-functional miRNPs with the increase in total cellular miRNP levels observed in Aβ_1-42_ oligomer-treated cells, C6 glioma cells were transfected with Renilla Luciferase (RL) reporter having 3’-UTR of IRAK1 mRNA with three miR-146a binding sites (Figure 3A). A reduction in miR-146a activity was observed in Aβ_1-42_ treated cells compared to the DMSO treated control cells (Figure 3B). The most-probable cause of reduced miRNA activity could be the unbinding of miRNA with Ago2 proteins that are known to happen in mammalian macrophage during its activation phase^19^ and also in neuronal cells during the nerve growth factor induced differentiation process^26^. To check whether a similar effects on Ago2 by Aβ_1-42_ may cause the down regulation of miRNA-Ago2 binding in Aβ_1-42_ treated astrocyte cells, we cultured primary astrocytes from rat pups and treated them with DMSO or Aβ_1-42_ for 24h. Ago2 was immunoprecipitated (IP) followed by qPCR based measurement of bound miRNAs. The RNA amount was normalized with the quantity of Ago2 protein recovered during the IP process. The quantification revealed, in Aβ_1-42_ treated cells, there has been increased association of Ago2 with miR-146a and miR-155 compared to the control cells (Figure 3C). We also found an increase in both Ago2 associated miR-16 and miR-29a level upon Aβ_1-42_ treatment (Figure S2A). We did similar experiments with primary neurons cultured *ex vivo*, where we could not detect a significant effect of Aβ_1-42_ oligomers on neuronal miRNA levels or Ago2 associated miRNA level suggesting a astrocyte specific effect of Aβ_1-42_ on cellular miRNPs (Figure 3D–3E). To check the possible role of any post translational modification affecting miRNA activity in Aβ-treatment context, we measured phosphorylated Ago2 levels in Aβ_1-42_ treated cells. Reduced phosphorylated Ago2 level in Aβ_1-42_ treated cells was detected compared to the control cells and the results correlated with reduced interaction between Ago2 and mTOR after Aβ_1-42_ treatment (Figure 3F–3G) indicating a possible role of mTOR in miRNA activity regulation via controlling Ago-phosphorylation. It was suggestive from the earlier experiments that Aβ_1-42_ mediated decrease in Ago2 phosphorylation might hinder dissociation of miRNA from Ago2 protein but reducing miRNA activity. So we hypothesized that lack of Ago2 phosphorylation might perturb the repression of target mRNA in mammalian cells. Ago2 proteins have a MID domain which is found to be important for binding with the 5’ phosphate group of small RNA^28^. A phosphorylation at the Y529 position in MID domain imparts a negative charge which dissociates miRNA from Ago2 protein. Ago2Y529F mutant that cannot be phosphorylated at Tyr529 position remains bound to miRNA^19, 28^. C6 glioblastoma cells were transiently transfected with the Ago2-WT and Ago2-Y529F protein in absence and presence of Aβ_1-42_ oligomers and qPCR was done to check the cellular mRNA levels of IL-1β and IL-6. We found a higher expression of both IL-1β and IL-6 mRNA level in Ago2-Y529F expressing cells even without Aβ_1-42_ exposure, which suggests the importance of Ago2 phosphorylation in cytokine mRNA expression (Figure 3H). However the effect of Aβ_1-42_ on cytokine expression was absent in cells expressing Ago2 Y529E. This further suggests that cytokine expression regulation primarily via Ago2 phophorylation in glial cells. To confirm the involvement of mTORC1 in miRNA activity regulation process possibly via Ago2 phosphorylation, cells were exposed to Rapamycin, a known mTORC1 blocker ^43^. We found significant decrease in miR-146a activity when treated with rapamycin (Figure 3I). We also found reduced amount of Ago2 phosphorylation and an increase in Ago2 bound miR-146a level upon rapamycin treatment (Figure 3J–3K). These data signifies the effect of mTORC1 inhibition in controlling miRNA activity and Ago2 phosphorylation that are also observed in astrocytes exposed to Aβ_1-42_. Experiments were also done in heterogeneous context using HEK293 cells depleted for Raptor an integral protein of mTORC1 complex^44^ and expressing exogenously a liver specific miR-122 there. This was done to recheck the interlinking of mTOR inactivation with Ago2 phosphorylation and miRNA activity regulation even in non-astroglial cells. In HEK293 cells treated with siRaptor, we documented a reduced miRNA activity and like in Aβ_1-42_ treated cells, accompanied by a reduction in Ago2-phosphorylation confirming mTORC1 involvement in miRNA activity regulation in general in mammalian cells by targeting Ago2 phosphorylation (Figure S2B-2C).

**Figure 3.**
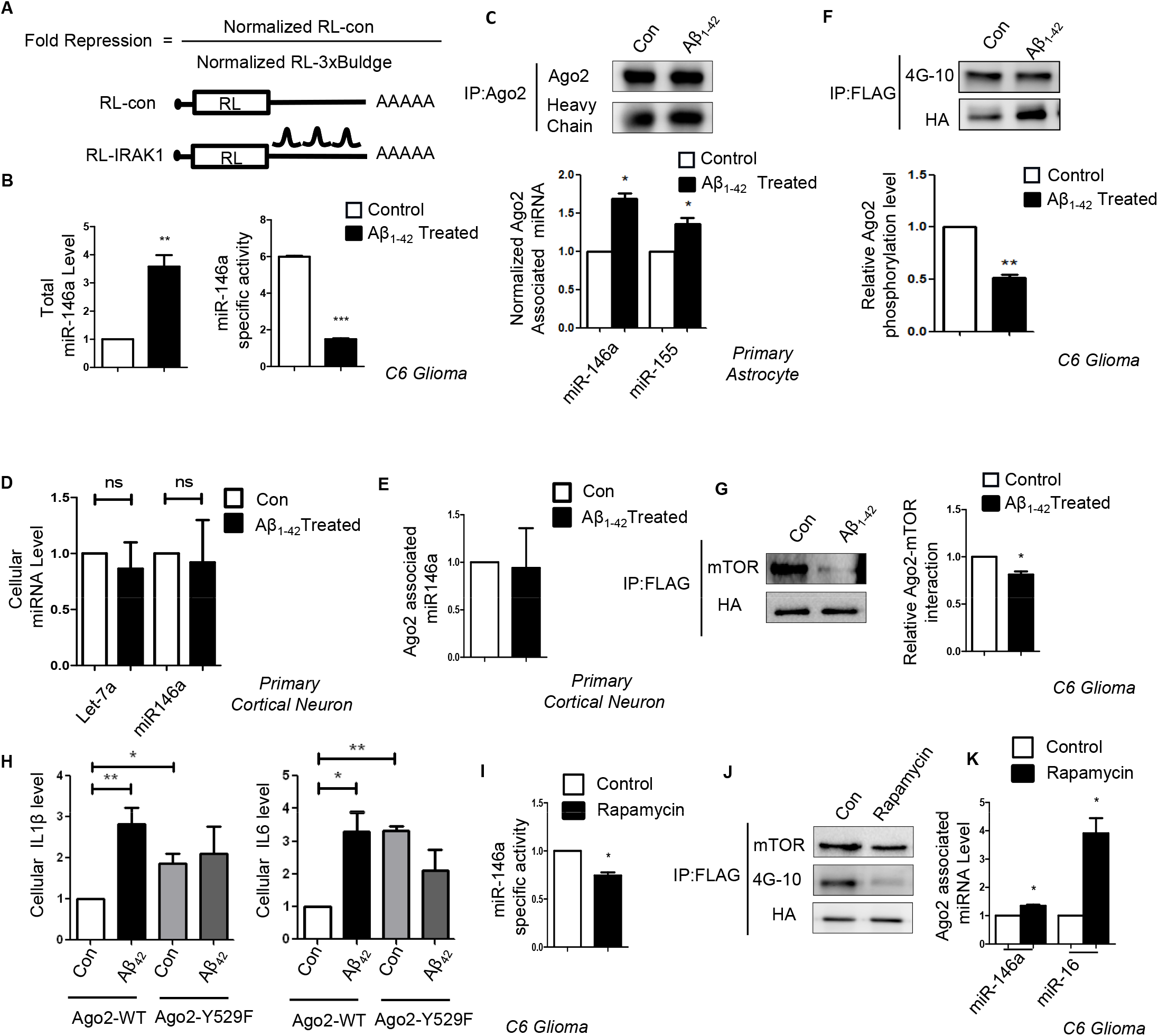
mTOR mediated Ago2 phosphorylation is required for miRNA activity. **A** Schematic design of different Renilla reporters (RL) used for scoring miRNA activity. 3’ UTR of IRAK1 mRNA with three miR-146a binding sites were cloned on 3’UTR of the Renilla mRNA. **B** Effect of Aβ_1-42_ treatment on cellular miR-146a activity and level. Specific activity of miR-146a was calculated by normalizing the fold repression of target reporter mRNA against the corresponding miRNA expression level. Increased cellular miR-146a level in C6 Glioblastoma cells upon Aβ_1-42_ treatment. The data was obtained by qRT-PCR based quantification and was normalized against the expression of U6 SnRNA. **C** Increase in Ago2-miRNA association in rat primary astrocytes upon Aβ_1-42_ treatment. Endogenous Ago2 from control and 1.5 μM Aβ_1-42_ treated primary rat astrocytes were immunoprecipitated and the levels of Ago2 associated miR-146a and miR-155 were estimated by qRT-PCR. Values were normalized against the amount of Ago2 obtained in each immunoprecipitation reaction. **D-E** No change in miRNP levels in rat cortical neurons exposed to 1.5μM Aβ_1-42_. Primary neuronal cells were treated with Aβ_1-42_ oligomers for 24h and amount of endogenous miRNA and miRNA associated with immunoprecipitated Ago2 were estimated and plotted after normalization against U6 RNA and Ago2 levels respectively. **F-G** Reduced mTOR-Ago2 interaction and Ago2 phosphorylation in Aβ_1-42_ oligomer treated C6 Glioblastoma cells. Western blot data showing the amount of Ago2 phosphorylation (F) and amount of mTOR pulled down with Ago2 (G). C6 Glioblastoma cells were transiently transfected with FH-Ago2 and then treated with 2.5μM of Aβ_1-42_ for 24h. Immunoseparation of FH-Ago2 was done with FLAG beads. The amount of Ago2 pulled down was detected by anti-HA antibody and levels of phosphorylated Ago2 (Tyr phosphorylation) was detected by anti p-Tyr antibody 4G10. **H** Effect of expression of Y529F mutant (Phospho mimetic mutant) of Ago2 on cellular IL-1β and IL-6 mRNA level on C6 cells. The qPCR data was normalized against GAPDH mRNA level. **I** Graph representing miR-146a specific activity upon rapamycin (100nM) treatment in C6 glioblastoma cells. The fold repression of miR-146a was normalized with the miR-146a expression level. **J** Western blot data showing level of Ago2 phosphorylation and Ago2-mTOR interaction upon rapamycin (100nM) treatment. **K** qPCR data showing Ago2 associated miR-146a and miR-16 level upon rapamycin (100nM) treatment. qPCR data was normalized with the amount of Ago2 pulled down from the immunoprecipitation reaction. For statistical significance, minimum three independent experiments were considered in each case unless otherwise mentioned and error bars are represented as mean ± S.E.M. P-values were calculated by utilizing Student’s t-test. ns: non-significant, *P < 0.05, **P < 0.01, ***P < 0.0001.

### Sequestration mTORC1 to early endosome causes miRNA inactivation and increased miRNP formation in glial cells

We found decrease in phophorylated S6K along with active phosphorylated mTOR levels in Aβ_1-42_ treated astrocyte cells (Figure 4A–4B). The expression of mTOR is tightly regulated in mammalian cells where different nutrients and growth factor like glucose, amino acid and insulin can activate mTORC1 depending upon the metabolic state of the cell^45^. Interestingly, it’s not only the presence of upstream signals, but mTORC1 also shows localization based activation. It is being reported earlier, that translocation of mTOR to the lysosomal membrane is important for mTORC1 activation where it can be activated by the GTPase Rheb ^34^. So to check for the possibility that Aβ_1-42_ mediated mTORC1 deactivation may happen because of the reduced mTORC1 localization to the lysosomal membrane, we microscopically looked at the localization of mTOR in control and treated cell. Confocal images were taken from both control and Aβ_1-42_ treated C6 glioblastoma cells, where lysosomes were stained with lysotracker (red) and mTOR (green) was visualized with indirect immunofluorescence (Figure 4C). There was a decrease in colocalization between mTOR and lysosome indicating a faulty translocation of mTOR in amyloid treated cells (Figure 4D). Since mTOR was not translocating at the lysosomal surface, it was important to find out the altered localization of mTOR upon Aβ_1-42_ exposure. We found a significant increase in colocalization between mTOR and early endosome (Figure 4E–4F). Cells were transiently transfected with YFP-Endo which marks early endosomes and then activated with oligomeric Aβ_1-42_ for 24hrs. Higher colocalization between early endosome and mTOR suggests Aβ_1-42_ restricts mTOR to the early endosomes which retards its translocation to the lysosomal surfaces to get activated. To check whether mTOR sequestration in early endosome can affect miRNA activity or miRNP formation, we expressed a constitutively active GTPase deficient Rab5 mutant, Rab5Q79C (Rab5-CA) that is known to cause mTOR sequestration in early endosome^46^ (Figure 4G–4H). With the over expression of Rab5-CA mutant, we observed a decrease in specific activity of let-7a (Figure 4I) and an increase in both cellular and Ago2-associated miR-146a (Figure 4J–4K). These results connect the mTOR compartmentalization defects in Aβ_1-42_ treated cells to functional inactivation of increased miRNPs observed.

**Figure 4.**
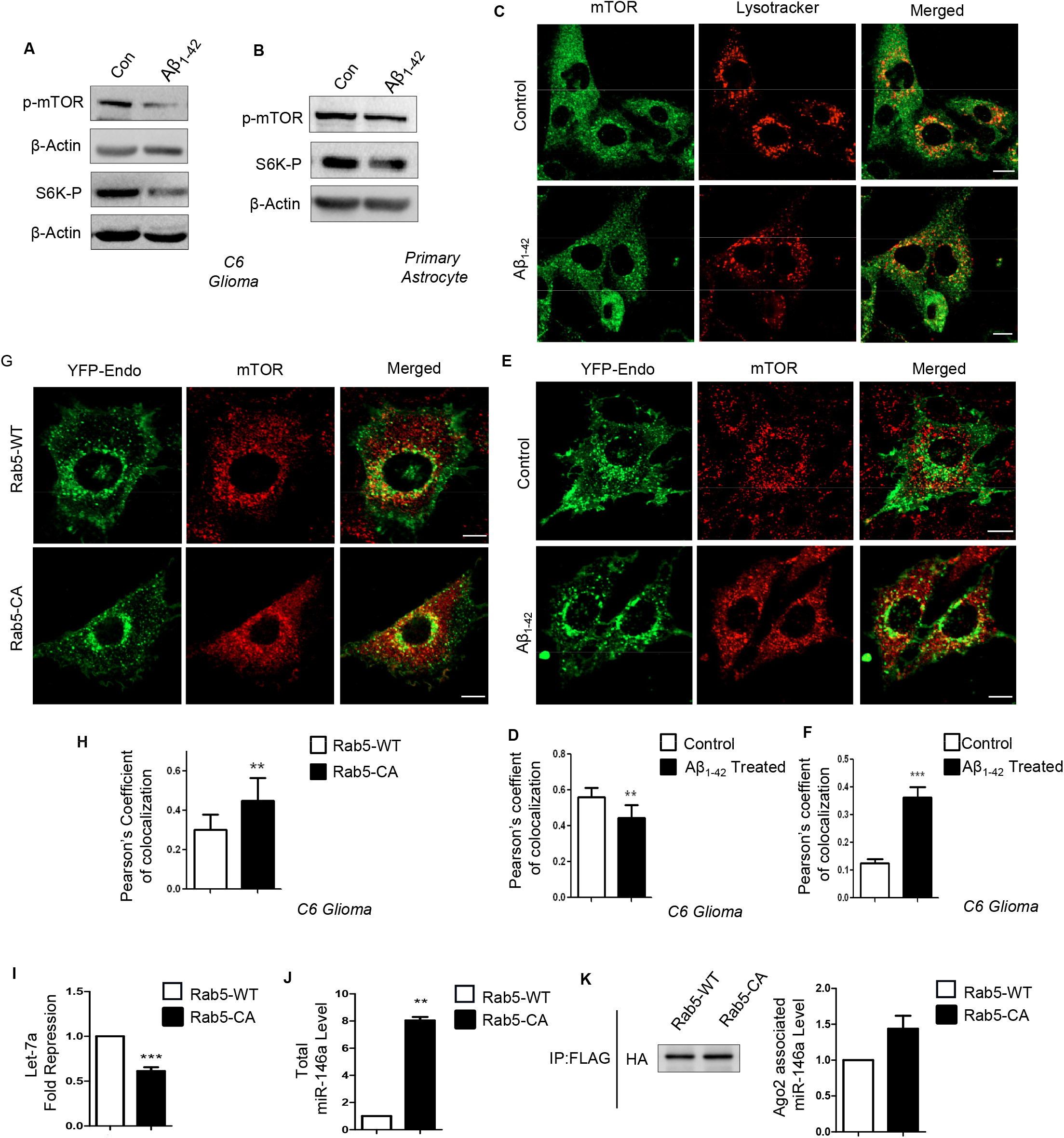
Sequestration to Endosome causes mTORC1 inactivation linked with increased miRNA-Ago2 interaction in glial cells. **A-B** Lowering of p-mTOR in Aβ_1-42_ oligomer treated C6 glioblastoma cells. Western blot data showing the levels of cellular p-mTOR level along with downstream substrate of mTOR phospho-S6K levels in C6 Glioblastoma cells (A) and in primary astrocyte cells (B) treated with Aβ_1-42_. **C-D** Decreased lysosome targeting of mTOR after Aβ_1-42_ oligomer treatment. Confocal image depicting localization of mTOR to lysosome in control and treated cells are shown in panel (C). Endogenous mTOR is visualized indirect immunefluorescence (green) and lysosome is labelled with lysotracker (red) in C6 Glioblastoma cells. Pearson’s coefficient of colocalization was used to measure the amount of mTOR translocating to lysosome in control and Aβ_1-42_ treated C6 Glioblastoma cells (D). **E-F** Increased endosome-mTOR localization after Aβ_1-42_ oligomer treatment. Confocal images showing localization of endosomes and mTOR. Endosomes were tagged with YFP-Endo (green) and endogenous mTOR (red) was detected by indirect immunofluorescence. Colocalization between endosomes and mTOR was visualized as yellow (E). Pearson’s coefficient of colocalization showing between early endosomes and mTOR in Aβ_1-42_ oligomer treated C6 Glioblastoma cells is shown in panel (F) **G-H** Effect of Rab5-CA on mTOR localization in C6 Glioblastoma cells. Colocalization of mTOR in C6 cells expressing YFP-Endo upon Rab5-CA expression was done. The cells with YFP-Endo (green) and mTOR (red) were visualized in control and Rab5-CA expressing cells (G). Pearson’s coefficient of colocalization of mTOR and Rab5-CA has been shown in panel (H). **I** Alteration of miRNA activity and levels in mammalian cells defective for endosome maturation due to expression of the constitutively active form of Rab5 protein. Drop of cellular activity of let-7a miRNA in C6 cells upon expression of constitutively activated Rab5 mutant Rab5-CA. The RL reporter with three let-7a miRNA binding sites was used to score the effect of Rab5-CA expression on miRNA repressive activity. **J-K** Levels of cellular and Ago2 associated miR-146a upon Rab5-CA expression in C6 Glioblastoma cells. The cellular miR-146 level was normalized against U6 snRNA level (J). The amount of Ago2 immunoprecipitated from control and Rab5-CA expressing cells were used for normalization of amount of miRNAs associated with Ago2 (K). For statistical significance, minimum three independent experiments were considered in each case unless otherwise mentioned and error bars are represented as mean ± S.E.M. P-values were calculated by utilising Student’s t-test. ns: non-significant, *P < 0.05, **P < 0.01, ***P < 0.0001.

### mTORC1 activation rescue cytokine repression by targeting miRNP reactivation pathway

Based on the results described so far, mTOR translocation to lysosome could be important for Ago2 phosphorylation leading to sustained miRNP recycling and activity in astrocyte cell. To confirm this mode of action of mTOR in glial cell, C6 cells were transfected with plasmids encoding Rheb-Myc before the Aβ_1-42_ oligomer treatment. Rheb is a constitutive activator of mTOR which can activate mTOR independently of growth factor or amino acid stimulation ^47, 48^ Results from Rheb expressing cells suggests rescue of mTOR localization upon Aβ_1-42_ treatment in Rheb-Myc expressing cells (Figure 5A). We observed about a 1.6 fold reduction in mTOR sequestration in endosome suggesting there was mobilization of mTOR from endosomal compartments (Figure 5A-B). To investigate a possible role of mTOR reactivation in miRNA activity, we checked the status of Ago2 positive bodies in Aβ_1-42_ treated cells expressing Rheb-Myc (Figure 5C). Interestingly in Rheb expressing cells, we could find a lesser number of Ago2 positive bodies colocalized with RNA processing bodies or PBs compared to that in Aβ_1-42_ treated cells (Figure 5D). These data clearly suggest a role of mTOR signaling in mobilizing the Ago2 from an otherwise inactive compartment into an active compartment where they could repress the target cytokines. Activation of mTOR activity upon Rheb-Myc expression was confirmed by measuring the levels of phosphorylated mTOR and S6 Kinase level in untreated and Aβ_1-42_ treated cells pre-transfected with control or Rheb-Myc expression plasmids (Figure 5E). To get further evidence whether this process restores miRNA activity *per se*, we checked the different cytokine levels. Quantitative estimation revealed that there has been a 2.7 fold decrease in IL-1β level and about 1.5 fold decrease in IL-6 level compared to the Aβ_1-42_ treated cells upon Rheb-Myc expression (Figure 5F). Again Rheb-Myc expressing cells showed a decrease in cellular miR-146a and miR-155 levels compared to the only Aβ_1-42_ treated cells (Figure 5G). Rheb mediated mTOR reactivation also increases Ago2 phosphorylation in Aβ_1-42_ treated cells reducing the Ago2 associated miR-146a and miR-155 level (Figure 5H and 5I). We also observed a similar effect in cellular and Ago2 associated miR-16 and miR-29a level when cells were expressing Rheb-Myc (Figure S3A-3C). Since miR-16 and miR-29a do not directly regulate expression of any proinflammatory cytokines, so there is a possibility that Aβ_1-42_ mediated mTOR inactivation mechanism may affect the function of quite a few miRNAs.

**Figure 5.**
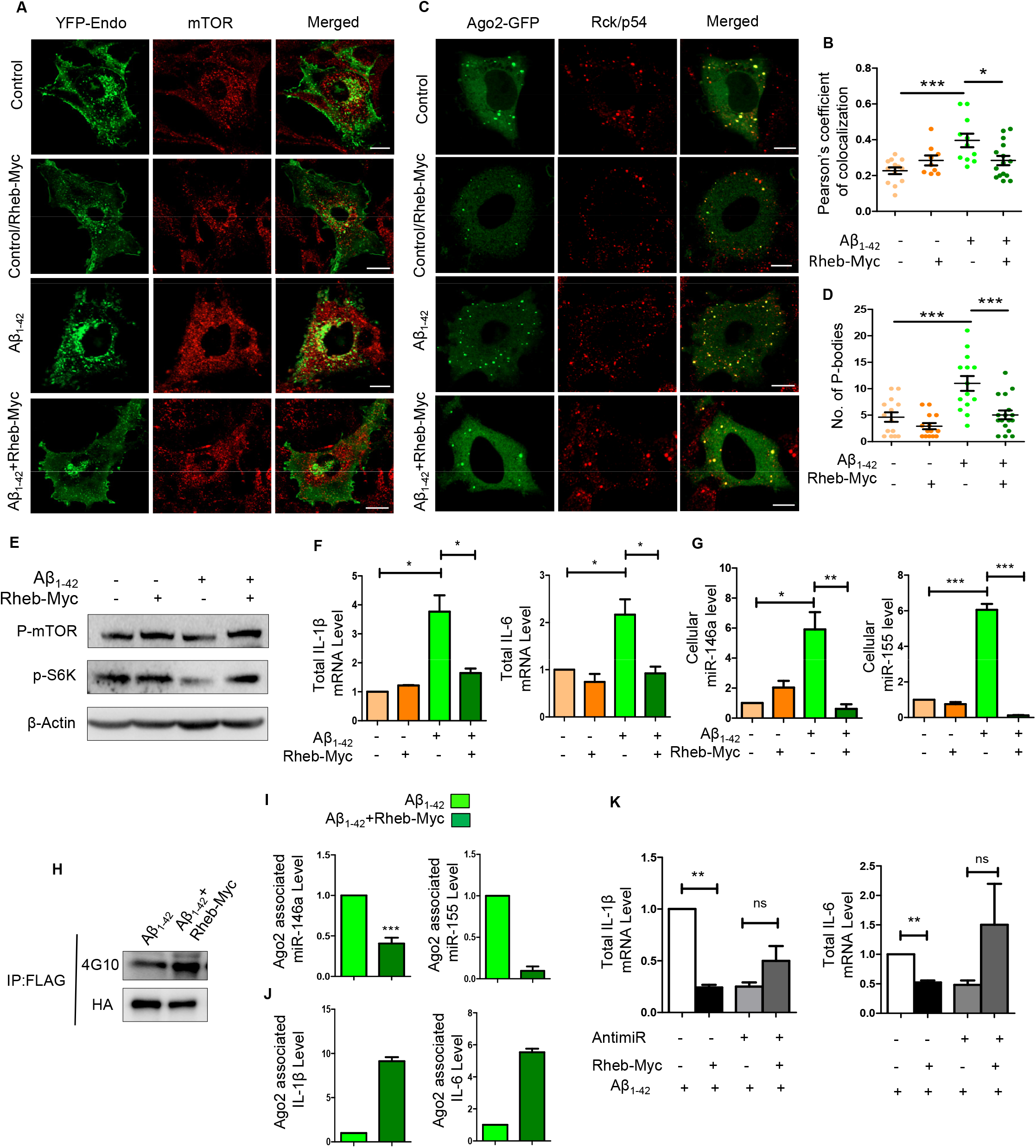
Reactivation of mTOR by Rheb-Myc mobilizes mTOR from endosomes and reduces cytokine production by targeting miRNA pathway. **A-B** Altered mTOR localization upon expression of Rheb-Myc in Aβ_1-42_ oligomer treated C6 Glioblastoma cells. Confocal images showing colocalization of endosomes and mTOR. The C6 Glioblastoma cells expressing YFP-Endo and thus having endosomes with tagged with YFP-Endo (green). Endogenous mTOR is stained in by indirect immunefluorescence (red). Yellow is used to show the extent of colocalization between endosomes and mTOR (A). Plot depicting amount of colocalization between mTOR and endosome in control and Aβ_1-42_ oligomer treated C6 glioblastoma cells transfected rather with control vector or with Rheb-Myc expression plasmid. Amount of colocalization is measured with Pearson’s coefficient of colocalization (B). **C-D** Ago2 localization to PBs gets affected in cells expressing Rheb-Myc. Confocal microscopic images of Ago2 bodies stained with GFP-Ago2 (green) co-stained for endogenous Rck/p54 (red). Colocalized Ago2 and Rck/p54 bodies were considered as PBs (yellow) (C). Plots depicting number of PBs per cell in in C6 glioblastoma cells that were untreated or treated with Aβ_1-42_. In both conditions effect of expression of Rheb-Myc was scored and amount of colocalization is measured with Pearson’s coefficient of colocalization (D) **E** Effect of Rheb-Myc expression on mTOR activation in control and Aβ_1-42_ oligomer treated C6 Glioblastom cells. Western blot data showing the levels of cellular p-mTOR and phospho S6K levels in untreated and cells expressing either control or Rheb-Myc expression vectors. **F-G** Expression of Rheb-Myc restores the cytokine and miRNA expression to control untreated levels in Aβ_1-42_ oligomer exposed C6 Glioblastoma cells. Quantitative data showing the effect of Rheb-Myc expression on cellular IL-1β and IL-6 mRNA level in both control and Aβ_1-42_ treated C6 Glioblastoma cells. The cytokine mRNA levels were normalized against GAPDH mRNA level (F). Effect of Rheb-Myc expression on cellular miR-146a and miR-155 levels in both control and Aβ_1-42_ treated C6 Glioblastoma cells (G). The data was normalized against U6 snRNA level. **H-J** Reversal of Ago2-miRNA and Ago2-cytokine mRNA interaction upon Rheb-Myc expression. Ago2 associated miR-146a and miR-155 levels and IL-1β and IL-6 mRNA levels were measured in C6 Glioblastoma cells pre-treated with Aβ_1-42_ oligomers (I and J). RNA level was normalized by the amount of Ago2 pulled down in immunoprecipation reaction (H). **K** Effect of Rheb-Myc expression on IL-1β and IL-6 mRNA level when the cells were pre-treated either with control or by miR-146a and miR-155 specific antagomiR. In all the sets C6 Glioblastoma cells were activated with 2.5μM of Aβ_1-42_ after 48h of Rheb-Myc and Anti-miR transfection. The data was normalized against GAPDH mRNA level. For statistical significance, minimum three independent experiments were considered in each case unless otherwise mentioned and error bars are represented as mean ± S.E.M. P-values were calculated by utilizing Student’s t-test. ns: non-significant, *P < 0.05, **P < 0.01, ***P < 0.0001.

Rheb-Myc expression has also increased the fraction of IL-1 β and IL-6 mRNA bound to Ago2 in Aβ_1-42_ treated cells-confirming availability of active miRNPs for repression of target messages (Figure 5J). miR-146 and miR-155 are the two major miRNAs reported as primary controller of cytokine expression and by trying the inactivation of both of these miRNAs with inhibitory antisense oligos, we can expect to find the reduced response of Rheb-Myc expression in Aβ_1-42_ treated cells. We found a drop in the IL-β level when Aβ_1-42_ exposed cells expressing Rheb-Myc (Fig. 5F). Now, we wanted to check whether Rheb-Myc mediated rescue in the cytokine mRNA repression depends on the repressive activity the miRNA. In the experiment done in Figure 5K all the C6 glioblastoma cells were treated with Aβ_1-42_. Cells expressing control vector, control Anti-miR-122 and exposed to 2.5μM Aβ_1-42_ were taken as control set. Cellular cytokine mRNA levels of control set were taken as 1 and rest of the experimental samples were plotted against it. Supporting the notion that the response of Rheb-Myc expression on cytokine expression is through reactivation of miR-155 and miR-146 in Aβ_1-42_ exposed cells, we have noted no significant reduction in cytokine production upon Rheb-Myc expression when the same cells were transfected with anti–miR-146 and anti-miR-155 oligos in combination (Figure 5K). This is expected as, due to lack of functional miR-146 and miR-155, the cytokines become non-responsive to mTOR activation and thus mTOR must affect primarily through these miRNAs on expression of the cytokines in glial cells. These results suggest activation of mTOR signaling to cause miRNP activity restoration in Aβ_1-42_ treated glial cells.

### Rescue of cognitive function in Aβ_1-42_ oligomer infused adult rat brain upon Rheb-Myc expression

Inactivation of mTORC1 pathway has been reported in AD brain^32, 49^. In the *in vitro* cell culture, we have documented restoration of mTORC1 activation to rescue cytokine repression by miRNPs. We checked whether over expression of Rheb-Myc in an Aβ_1-42_ oligomer induced AD model of the disease can result in reduction of pro-inflammatory cytokine expression to cause improvement in cognitive function of treated animal against the control group of animals. It has been shown before that Aβ_1-42_ oligomer infusion in the brain resulted in neuron death and deposition of Aβ plaque in the vicinity of infusion site with defective cognitive functions ^39^. To check the effect of Rheb-Myc on Aβ_1-42_ induced neuroinflammation and cognitive functions, we injected Aβ_1-42_ oligomers along with Rheb-Myc expression cassette or control vector bilaterally in the hippocampal area of the brain of adult rat and kept for 15 days. Following this infusion, several memory function tests were performed to assess the cognitive functions in experimental group of animals (Figure 6A).

**Figure 6.**
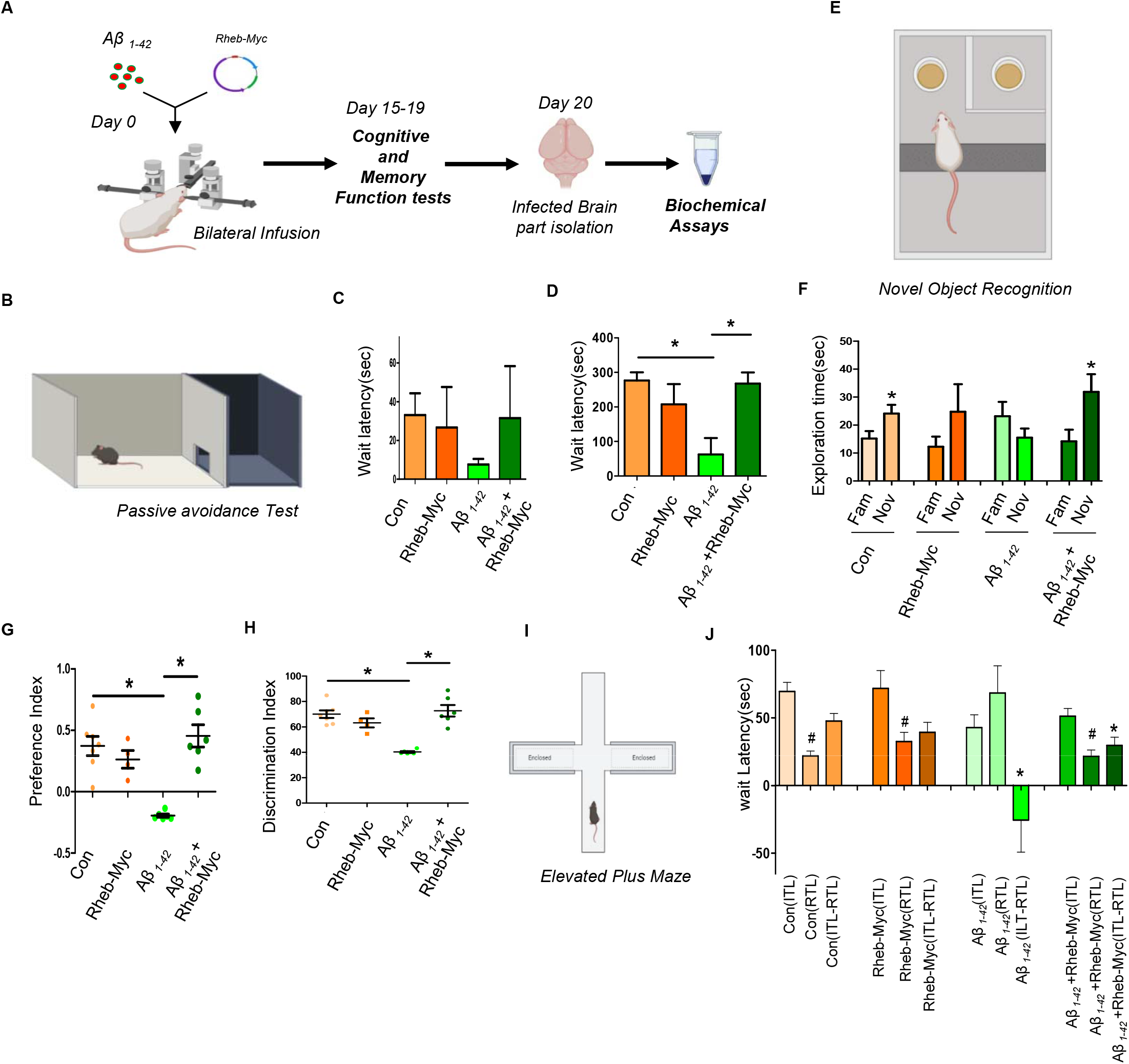
Rescue of cognitive function in Aβ_1-42_ infused adult rat brain upon Rheb-Myc expression. **A** Experimental flow chart showing a bilateral infusion of Aβ_1-42_ oligomers and Rheb-Myc plasmid was done in adult rat hippocampus. After 15 days of incubation, different behavioral assays were carried out followed by sacrificing the animals. **B-D** Rescue of wait latency in Aβ_1-42_ infused adult rat brain upon Rheb-Myc expression. Passive avoidance assessment of memory performance was done and the graph shows the acquisition process of experimental subjects. The rats that are being able to enter the dark chamber within 300 seconds were selected as eligible for testing. The bar diagram reveals the wait latency in different experimental sets of rats. Rats having Rheb-Myc plasmid infused along with Aβ_1-42_, showed higher wait latency on the probing day, as compared to the animals having only Aβ_1-42_ infusion(D) (n=7). **E-H** Novel object recognition test to show that the short term memory recognition capability has been restored by Rheb-Myc expression in Aβ_1-42_ injected animals. Aβ_1-42_ and Rheb-Myc expression plasmid co-infused animals showed improvement in novel object identification capacity as compared to the Aβ_1-42_ infused rodents (F). The Rheb-Myc-Aβ_1-42_ co-infused animals also showed increased preferential index and discrimination index as compared to the Aβ_1-42_ infused group of animals indicating improvement in learning and memory function (G and H)(n=7). **I-J** The Elevated Plus Maze analysis showed improvement in cognitive ability in Rheb-Myc expressing animals. The retention transfer latency (RTL) was measured in all control and Aβ_1-42_ infused groups and the difference of Initial Transfer Latency (ITL) and RTL was plotted along with the ITL and RTL separately (n=7). For statistical significance, minimum three independent experiments were considered in each case unless otherwise mentioned. The P-values were obtained with the help of Non-parametric Student’s t-test. Animal data are represented as means +SEM and analyzed by one way ANOVA followed by Bonferroni’s test.

We have performed the passive avoidance test which is used to understand learning and memory defects of rodents. This test showed that intra-hippocampal Aβ_1-42_ infusion significantly reduced wait latency compared to control group of animals. Wait latency represents the time each animal spends in the light chamber before entering into the dark chamber where they experience a shock upon entering (experimental detail is described in the methods section) during the 300 sec period following 5 min habituation (Figure 6B). The probe test was performed 24h after the training test. Compared to the control group, injection of Aβ_1-42_ markedly reduced the retention of fear memory as they showed lower wait latency on probe test as they moved to dark chamber within few seconds. However, rats having Rheb-Myc plasmid infused in the hippocampus along with Aβ_1-42_ oligomer showed higher wait latency on probe stage as compared to the animals having only Aβ_1-42_ infused (Figure 6 C-D). After behavioral studies animals were sacrificed. Brain tissues were fixed and immunohistochemistry was done to check the expression of both Rheb-Myc and ZS-Green (ZS-Green encoding plasmid was co-injected with Rheb-Myc encoding plasmid) at infusion site. We could find colocalization between both ZS-Green and Rheb-Myc at the infection site confirming their expression (Figure S4A-4B).This result indicates that Rheb-Myc could reverse the declining cognitive function in Aβ_1-42_ infused rats.

Next we performed the novel object recognition test and observed that Aβ_1-42_ infused animals had no preference towards the novel object as compared to control group. In contrast, this short term memory recognition capability in Aβ_1-42_ co-infused animals was significantly improved upon Rheb-Myc expression. The Rheb-Myc-Aβ infused animals also showed increased preferential index and discrimination index as compared to the Aβ_1-42_ infused group of animals (Figure 6 E-H). Additionally, the elevated plus maze (EPM) analysis showed improvement in cognitive ability in the above-mentioned group of animals. In Figure 6I, latency was the measure of the time taken by the animal to move from the open to the closed arm in EPM. In the Aβ-infused group, retention transfer latency (RTL) was higher than initial transfer latency (ITL) which was significantly reduced in other groups. The difference between ITL and RTL (ITL-RTL) was used to assess memory and learning in EPM test. Notably, a negative difference was obtained in the Aβ_1-42_ infused animals (Figure 6I). However, Aβ_1-42_ infused rats injected with Rheb-Myc plasmid showed positive difference between ITL and RTL compared to Aβ-infused group (Figure 6I-J).

The immunohistochemistry of Aβ_1-42_ infused adult rat brains showed astrogliosis depicting increase in GFAP in astrocyte cells but in Rheb-Myc plasmid and Aβ coinfused rat brain, enhanced GFAP was found to decrease as compared to the Aβ_1-42_ oligomer infused rat brain. However no decrease in Aβ accumulation was found in Rheb expressed Aβ_1-42_ infused brains that suggest the effect of Rheb-Myc was on glial cell response and cytokine production rather than an effect on Aβ deposition (Figure 7A-B). These results indicate that ectopic expression of Rheb protein in Aβ_1-42_ infused rat brain causes improvement in cognitive behavior and spatial memory. While checking the differential expression status of cytokine mRNA levels in the infused rat brains, almost 2 folds decrease in proinflammatory IL-6, IL-1β was observed in the Rheb-Myc-Aβ_1-42_ co-infused rats as compared to the Aβ_1-42_ only infused rats (Figure 7C-D). Rheb-Myc expressed animals show a decrease in the both cellular and Ago2 associated miR-146a level compared to the only Aβ_1-42_ injected animal tissue (Figure 7E-F). Therefore by activating mTOR and miRNP-recycling the cytokine expression could be controlled to rescue the memory loss and related phenotype in AD model of Alzheimer’s disease by expressing Rheb (Figure 7G).

**Figure 7.**
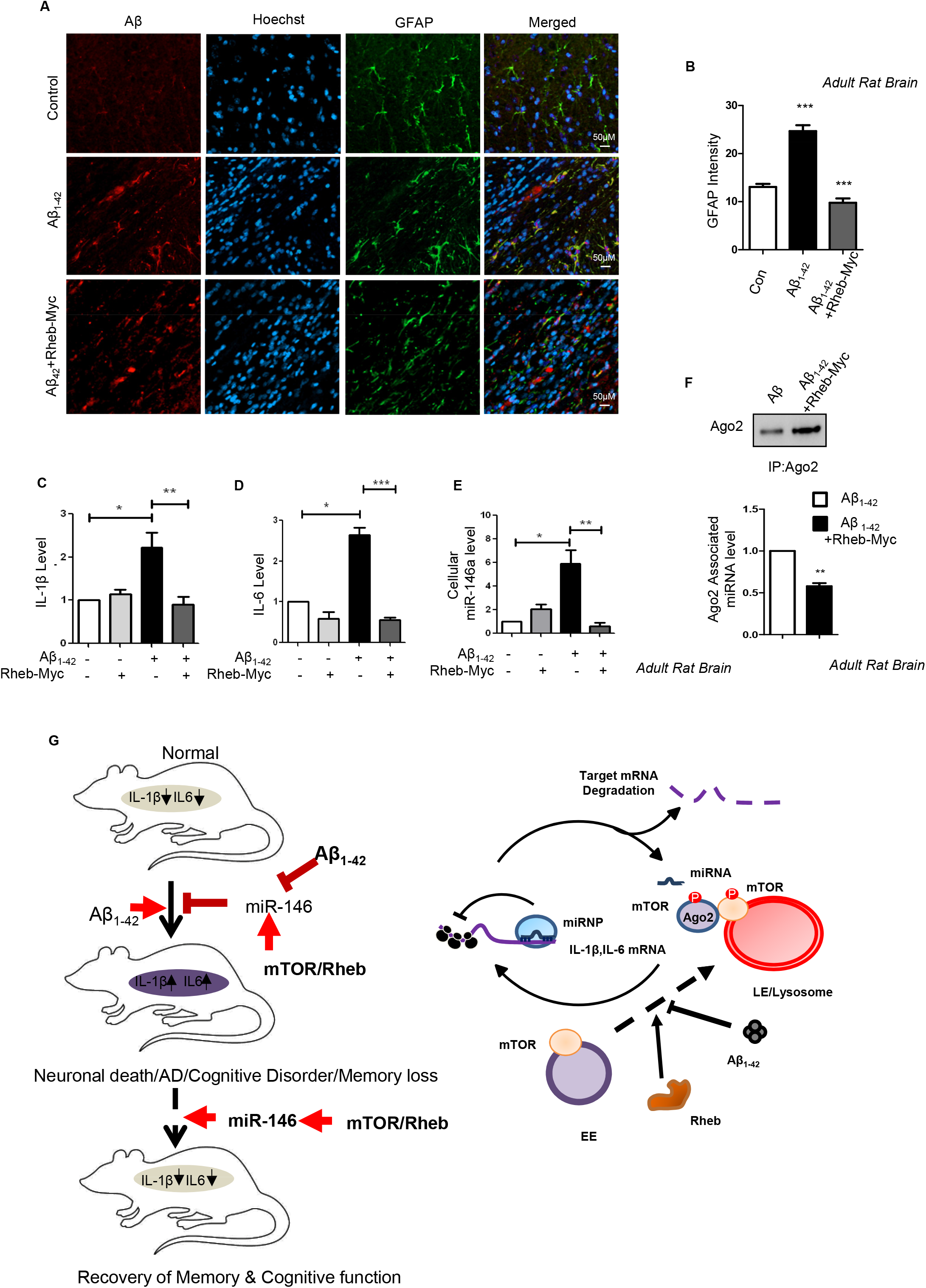
Rescue of cytokine level in Aβ_1-42_ infused adult rat brain upon Rheb-Myc expression. **A-B** Histology sections of Aβ_1-42_ infused and control rat brains. Cryosections, which were subjected to immunohistochemistry (IHC) are shown. Confocal images of the sections of Aβ_1-42_ infused adult rat brains showed astrogliosis depicted by increase in GFAP positive astrocyte cells. This was reversed in Rheb-Myc-Aβ infused rat brain, where enhanced GFAP was found to decrease as compared to the Aβ_1-42_ infused rat brain. (A). The graph depicting the intensity of GFAP in rat hippocampus is shown in panel (B). **C-D** Expression of IL-1β (C) and IL-6 mRNA (D) level in rat hippocampal tissue sample upon Rheb-Myc expression. The data was normalised against GAPDH mRNA level. **E** Cellular miR-146a level in the infused rat brains. The values were obtained by qRT-PCR and normalized against the U6 snRNA level. **F** Levels of Ago2 associated miR-146a level gets restored by Rheb-Myc in Aβ_1-42_ infused Rat hippocampus tissue. Immunoprecipitation was done for endogenous Ago2 and associated miRNA was measure with qRT-PCR. The value was normalized against Ago2 band intensity measured from the western blot. **G** The model depicts how Amyloid beta exposure causes mTOR entrapment with endosomes restricting miRNP target RNA interaction. This causes inactivation of miRNP to elevate proinflammatory cytokine production in glial cells. Interestingly ectopic activation of mTOR by Rheb restores miRNP function in diseased brain rescuing the cognitive function in adult rat brain. The mechanism of action of Aβ_1-42_ on miRNA via mTOR has been shown in right panel. For statistical significance, minimum three independent experiments were considered in each case unless otherwise mentioned and error bars are represented as mean ± S.E.M. P-values were calculated by utilising Student’s t-test. ns: non-significant, *P < 0.05, **P < 0.01, ***P < 0.0001.

## Discussion

Several miRNAs are expressed in brain and thought to regulate thousands of transcripts within the brain^3^. In the present analysis, we could identify de-regulated miRNA and mRNA within the cortical regions of AD patients which may be associated with AD development or progression. Interestingly, in our analysis, a substantial percentage of up-regulated miRNA also found to have a higher fraction of their corresponding target mRNAs as up-regulated. This trend was consistently observed in late stage Alzheimer’s disease cases considered from multiple studies. These observations suggest that in contrary to the general regulatory relationship between miRNA and their target mRNA wherein miRNA mainly down-regulate mRNA, other regulator to target relationships might also exist in tissues under disease conditions. In particular, miR-146a-5p is one such miRNA that exhibited this phenomenon wherein increased production of miR-146a-5p resulted in the up-regulation of multiple target mRNAs.

So any change in the activity of the miRNA pathway could play a significant role during development or during any pathological condition. Also molecular mechanism behind Aβ mediated neuronal death is still elusive and an interplay between cellular structure, molecular biological processes and resulted immune response of the disease could contribute to the fact that why AD is so difficult to treat. Another important aspect was to look at the involvement of glial cells in the buildup and disease progression as glial cells easily outnumber neuron in the brain^50^. So in the early stage of the disease, glial cells might be involved in rectifying the problems of the neurons due to Aβ_1-42_ oligomer deposition, but eventually contribute to the prognosis of the disease.

We found mTOR sequesters into early endosome with exposure to Aβ_1-42_ in glial cells. mTORC1 is a very important protein complex that can regulate the neuronal differentiation and brain activity. Depletion of an essential mTORC1 component Raptor is known to reduce the brain size considerably^51^. mTORC1 signaling can also play a role in phosphorylation of Ago2 protein which eventually can affect the miRNP function. Rheb, which is a GTPase and an activator of mTORC1, can reverse this process and results in decrease in Ago2 miRNP content. mTORC1 activity driven pathways may regulate miRNA maturation process as well. Tuberous Sclerosis Complex (TSC) is an autosomal dominant disorder and caused by mutations in Tsc1 or Tsc2 that is connected with mammalian target of rapamycin (mTOR) hyperactivity. TSC knockout cell which has an increased mTORC1 activity has reduced mature miRNA level^52^. So mTOR activation by Rheb can bring down the maturation process and also recycle of existing miRNPs and both these pathways may contribute in the miRNA activity rescue process. Rheb knockout animals are embryonic lethal ^53, 54^. Also conditional Rheb knockout adult mouse have limited life span, indicating Rheb is an essential protein factor which could play a major role in brain development and function. Here, injecting Rheb expressing plasmid into animal brain, we could improve motor function as well as memory of the AD animals. Similar to that, researchers have shown Rheb expression in brain of HD mouse model resulted in improve motor function^55^. Also in Parkinson’s Disease, Rheb expression in dopaminergic neurons promotes axonal regrowth^56^. Therefore, Rheb can affect brain function at multiple points. For example, mTOR mediated translation, a Rheb dependent process, is important at synapse for maintaining synaptic plasticity and at cell body for maintaining memory^57, 58^. Also, Rheb expression can increase acetyl choline level in the brain that can improve brain cognitive functions^59^. Although all these factors may play a role, our data suggests miRNA activity regained by Rheb expression may be the most important factor in recovery of behavioral, cognition and memory function in AD.

We have stereotactically injected Aβ_1-42_ oligomers in adult rat brain to produce the amyloid plaques in the brain. This model has several advantages as we can monitor the early cellular events during plaque formation and the effects it has on the miRNA machinery during early phase of the disease. Also oligomers can be taken up by glial cell in both the *in vivo* and *in vitro* cell culture system which can have additional physiological effects on the cellular structure and machinery. This allow the limited and spatio-temporal specific exposure of brain cells to Aβ_1-42_ and thus is advantageous over the transgenic mouse models having mutated Aβ gene (APP) or related protease gene which may have some compensatory effect on both structure and physiology in the adult brain. Additionally, the transgene is expressed at a very early stage and in all the cells of the brain. Importantly the number of AD cases reported with genetic mutations in APP gene is very limited compared to the higher number of cases noted with sporadic Aβ expression^60^. We have also injected Rheb expression vector with Aβ_1-42_ oligomer in brain hippocampus. This is to ensure only that portion of the brain which is exposed to Aβ_1-42_ oligomer only takes up the Rheb expressing plasmid and not the entire brain which could complicate the analysis. This method can be useful in introducing specific miRNA or antisense RNA in specific part of the brain with the help of co-injected Aβ_1-42_ oligomer.

Although we have only observed the effect Aβ_1-42_ oligomer has on miRNA machinery on astroglial cells, it would be interesting to see what effect it may have on the miRNAs in other glial cells. Glial cells outnumbers neurons in 10:1 ratio and among them almost 40% cells are astrocytes, making them one of the most abundant cells of the brain. Also Aβ_1-42_ oligomer injected brain tissues showed considerable astrocyte activation along Aβ deposition. Primary astrocyte shows elevated cytokine mRNA level when exposed to Aβ_1-42_ oligomer. We did find some changes in neuronal miRNA expression level but a drop in miRNA activity resulting in cytokine overproduction was found to be specific for astroglial cells and not for neurons. In LPS activated macrophage cells miRNAs play a significant role in reducing pro-inflammatory cytokines like TNF-α, IL-1β or IL-6 so that anti-inflammatory IL-10 production can initiate^19^. This is a very important step in inflammation process where miRNA fine tunes the balance between pro-and anti-inflammatory cytokine levels to maintain the homeostasis of the process. The astrocytes cells exposed to Aβ_1-42_ oligomer show a loss in miRNA activity where loss of Ago2 phosphorylation leads to building-up of inactive miRNP in astrocyte cells which restrict their encounter with the target mRNA. This leads to uncontrolled production of pro-inflammatory cytokine that eventually creates the neuroinflammation leading to loss of neurons.

If the mechanism is happening through Ago2 phosphorylation then this mechanism should affect other miRNAs as well. We found the pattern of data of miR-16 and 29a are very similar to what we also got for miR-146a and miR-155. Since miR-16 and miR-29a do not directly regulate expression of pro-inflammatory cytokines, we think there is a possibility that this mechanism affects the functions other miRNAs as well. In this context, it may be noted that in primary astrocytes and rat brain (also in AD patient brain data) we have documented increased expression of many miRNAs that are not even astrocyte specific or not a known regulator of cytokines. This also signifies the common pathway of miRNA activity modulation by mTORC1 has been targeted by Aβ_1-42_ oligomers. However, there are miRNAs that may be regulated at transcriptional level in a significant way, while there could be a subset of miRNAs that also have regulated export and turnover in Aβ_1-42_ treated cells and thus these factors may contribute significantly to the overall cellular levels of those miRNA to show a reduction in the overall expression level.

Our findings suggest, restoring mTOR activity in the glial cells can rescue disease phenotype and also cognition and memory in adult animals. Previous studies in the field of neurodegenerative diseases suggest maintenance of homeostasis of mTOR activity could be beneficial for the disease management. For example, in case of Autism, inactivation of mTOR is found to be beneficial as in autism patients hyper activation of mTOR is common. Again in ALS, where mTOR activity is reduced, restoring the activity found to be advantageous in disease prognosis^61, 62^. In a nutshell our study emphasis the complex circuit between metabolic pathways with gene expression regulation having an effect on disease pathology. An intervention with gene therapy or with small molecules to sustain normal miRNA metabolic pathways in brain will have a therapeutic value in a complex disease like AD.

## Materials and Methods

### Expression profile analysis of miRNA and their corresponding target mRNA in Alzheimer’s disease and cancer

Expression profiles of miRNA and mRNA were utilized to study the possible regulatory relationships between miRNA and their corresponding target mRNA in multiple cortical regions of Alzheimer’s disease patients as opposed to normal deceased individuals. With the help of R packages (R Core Team.,2017); ^63^ differentially expressed (fold change: +/-1.25 and p-value <= 0.05) miRNAs within the pre-frontal cortex and mRNA within different regions of the frontal cortex were determined in Alzheimer’s disease patients brain. The differentially expressed miRNA and mRNA identified in this manner considering patient samples are likely to be associated with the development or progression of Alzheimer’s disease [GEO datasets : GSE48552, GSE5281, GSE53697, GSE15222 ^64–69^]. Utilising miRNA-target regulatory information from Tarbase ^70^ and miRTarBase ^71^ databases, “regulator(s) to target(s)” (miRNA:mRNA) maps for the differentially expressed miRNA and mRNA were determined. Subsequently, we have compared the fractions of target mRNA of each differentially expressed miRNA that are significantly up-regulated or down-regulated. Additionally, to ascertain whether such “regulator(s) to target(s)” patterns could be occurring in other disease conditions, we have performed a disease enrichment analysis considering only the genes that were commonly up-regulated among the Alzheimer’s disease datasets considered ^72^.

In order to study “regulator(s) to target(s)” patterns in cancer, we have analyzed miRNA and mRNA expression profiling data from different cancer tissue samples. With the help of R packages, differentially expressed (fold change:+/-1.25 and p-value <= 0.05) miRNA and mRNA were determined in breast cancer, colorectal cancer and oral squamous cell carcinoma by considering relevant GEO datasets [GSE40056/GSE40057, GSE35982, GSE70664/GSE70665] ^63, 73–75^ R Core Team.,2017). Subsequently, the fractions of significantly up-regulated or down-regulated target mRNA of each of the differentially expressed miRNA under each condition were compared.

### SiRNA and Plasmid constructs

Plasmid information about pRL-con, pRL-3XBulge-Let7a, pRL-3XBulge-miR-122 and Firefly plasmids were previously explained in Pillai et. al 2005 and a kind gift from Witold Filipowicz. miR-122 was expressed from pmiR-122 plasmid which previously described ^76^. FLAG-HA-Ago2 and FLAG-HA-Ago2-Y529F were obtained as a kind gift from Gunter Meister. The plasmids Rab5-CA and Rheb-Myc were previously used ^77^, and were kind gift from J.M. Backer. All Si-RNAs were purchased from Dharmacon Inc. (On tatget plus Si-RNA).

### Cell Culture and reagents

HEK293 and C6 Glioblastoma cells were grown in Dulbecco’s modified Eagles medium (DMEM,Gibco) supplemented with 2mM L-glutamine,10% heat inactivated fetal bovine serum (FBS).

Primary astrocytes were cultured from 0 day old Sprague Dawley rat pups. The whole brain was dissected out and meninges were removed carefully. The neocortex parts were isolated and cut into pieces. The small cortical tissue pieces were minced and subjected to trypsinization for 30 min. Trypsinized brain tissue was triturated in DMEM (GIBCO) medium supplemented with 10% fetal bovine serum (FBS) (GIBCO) and passed through nylon mesh to avoid clumps. This single cell suspension was then added onto PDL (Sigma-Aldrich, St. Louis, MO, USA) coated plate and incubated for 2–3 min for preferential sticking of neurons. Next, the unattached cells were collected and harvested by centrifugation. Cells were resuspended in a fresh medium and seeded in a density of 1.2 million/35 mm plate or 0.4 million/well of a 24 well plate. Cells were maintained for 13 days in vitro (DIV) with a medium change given every alternate day.

Plasmid and Si-RNAs were transfected with Lipofectamine 2000 (Invitrogen) and RNAi max (Invitrogen) respectively following manufacture’s protocol. All the SiRNA and plasmid co-transfection was done using Lipofectamine 2000. For activation of Primary rat astrocytes or neurons 1.5uM of Aβ_1-42_ was used, whereas 2.5 uM of Aβ_1-42_ was used for activating C6 Glioblastoma cells. 100nM Rapamycin (Sigma) was used for blocking mTORC1.

### Preparation of Amyloid beta

HPLC-purified Aβ_1-42_ was purchased from American Peptide (Sunnyvale, CA, USA) and oligomeric Aβ_1-42_ was prepared as described previously ^39^. Briefly, lyophilized Aβ_1-42_ was reconstituted in 100% 1,1,1,3,3,3 hexafluoro-2-propanol (HFIP) to 1 mM. HFIP was removed by evaporation in a SpeedVac (Eppendorf, Hamburg, Germany), then resuspended in 5mM anhydrous DMSO. This stock was then stored at 80°C. The stock was diluted with PBS to a final concentration of 400 mM, and SDS was added to a final concentration of 0.2%. The resulting solution was incubated at 37°C for 18–24 h. The preparation was diluted again with PBS to a final concentration of 100 mM and incubated at 37°C for 18–24 h before use.

### Brain stereotaxic surgery

Male Sprague-Dawley rats (280–320g) were anesthetized by injecting thiopentone 50 mg/ kg (Thiosol sodium, Neon laboratories, Mumbai) and placing them on a steriotaxic frame (Stoelting, USA). Injection was done by using a 27-gauge Hamilton syringe. A volume of 5 μl of 100 μM Aβ in PBS and 3μg (4μl) of plasmid was infused at a flow rate of 0.5μl/min in the hippocampus at stereotaxic co-ordinates from bregma: AP: 3.6 mm, L: 2.1mm, DV: 2.8 mm, according to the rat brain atlas (Paxinos and Watson, 1982). An equal volume of PBS was injected in control animals. Proper postoperative care was taken to maintain proper health condition of the animal. Behavioral tests were performed fourteen days post stereotaxic surgery of the animal. Animals were sacrificed twenty days after following behavioral analysis. The brains were dissected out, following cardiac perfusion, and fixed in 4% PFA for 24h. They were then incubated in a 30% sucrose solution for another 24h before proceeding for cryo-sectioning. Cryo sections of the brain were done in cryotome (Thermo Shandon, Pittsburgh, PA, USA).

### Immunohistochemistry of brain slices

20 μm coronal cryo-sections of the brain from Aβ-infused or PBS-infused rats were blocked with 4 % BSA in PBS containing 0.43% Triton-X 100 for 1h-20min at room temperature. Brain slices were incubated in primary antibody in a blocking solution overnight at 4°C. Sections were washed thrice with PBS and incubated with a fluorescence-tagged secondary antibody for 2h at room temperature. Following three washes with PBS and Hoechst staining for the nucleus, the sections were mounted with DPX and observed in confocal microscope.

### RNA isolation and qPCR

Total RNA was isolated by using TriZol or TriZol LS reagent (Invitrogen) according to the manufacturer’s protocol. MiRNA assays by real time PCR was performed using specific Taqman primers (Invitrogen). U6 snRNA was used as an endogenous control. Real time analyses by two-step RT-PCR was performed for quantification of miRNA levels on Bio-Rad CFX96TM real time system using Applied Biosystems Taqman chemistry based miRNA assay system. One third of the reverse transcription mix was subjected to PCR amplification with TaqMan^®^ Universal PCR Master Mix No AmpErase (Applied Biosystems) and the respective TaqMan^®^ reagents for target miRNA. Samples were analyzed in triplicates. The comparative Ct method which included normalization by the U6 snRNA was used for relative quantification. For quantification of mRNA level, 200ng of total cellular RNA was subjected to cDNA preparation followed by qPCR by SYBR Green method. Each sample was analyzed in triplicates using comparative Ct method. Each mRNA levels were normalized with GAPDH as loading control.

### Immunoprecipitation assay

For immunoprecipitation of Argonaute2, cells were either transfected with FLAG-HA tagged Ago2 or endogenous Ago2 were pulled down from primary cells and tissue samples. For IP reactions, cells were lysed in Lysis buffer (20 mM Tris-HCl, pH 7.5, 150 mM KCl, 5 mM MgCl2, 1 mM DTT,10mM Sodium orthovanadate, 15mM NaF), 0.5% Triton X-100, 0.5% sodium deoxycholate and 1X EDTA-free protease inhibitor cocktail (Roche) for 30 min at 4□C, followed by three sonication pulses of 10 sec each. The lysates, clarified by centrifugation, were incubated with HA antibody (at a conc. of 1:100) pre-bound Protein-G Agarose bead (Invitrogen) or with pre-blocked anti-FLAG M2 beads and rotated overnight at 4□C. Subsequently, the beads were washed thrice with IP buffer (20mM Tris-HCl pH 7.5,150mM KCl, 5mM MgCl2, 1mM DTT,10mM Sodium orthovanadate, 15mM NaF). Washed beads were divided into two equal parts and each parts were analysed for bound proteins and RNAs by Western Blot and qPCR respectively.

### Luciferase Assay

miRNA repression was observed by doing dual luciferase assay. 10ng of RL-con and RL-3xB-Let-7a, RL-3xB-122 or RL-IRAK1 encoding plasmids was co-transfected with 100ng of Firefly (FF) encoding plasmid per well of a 24 well plate to study endogenous let-7a or exogenous miR-122 repression. Cells were lysed with 1X Passive lysis buffer (1X PLB Promega) before subjected to dual luciferase assay (Promega, Madison, WI) following the supplier’s protocol on a VICTOR X3 Plate Reader with injector (PerkinElmer, Waltham, MA). RL expression levels for control and reporter was normalized with the FF expression level for each reaction. All samples were done in triplicates.

### Immunoblotting

SDS-polyacrylamide gel electrophoresis was performed with the samples which include cell lysates, membrane fractions, immunoprecipitated proteins followed by transfer of the same to PVDF nylon membranes. Specific required antibodies were used to probe the blot for at least 16 hrs at 4°C. This antibody associated overnight incubation was followed by three washes with TBST (Tris buffer saline-tween 20), after which at room temperature, the membranes were incubated for 1h with horseradish peroxidase-conjugated secondary antibodies(1:8000 dilutions).Images of developed western blots were taken using an UVP BioImager 600system equipped with VisionWorks Life Science software (UVP) V6.80.

### Immunofluorescence and Confocal Imaging

For Immunofluorescence studies, primary and C6 Glioblastoma cells were grown on 18mm cover-slips coated with poly-D-lysine and gelatin respectively. Cells were transfected on the cover-slips as discussed previously and fixed for 30 mins in the dark with 4% paraformaldehyde solution after 48h. of transfection. Cells were blocked and permeabilized with 3% BSA containing 0.1% Triton X-100 for 30 min, followed by overnight incubation with specific primary antibodies at 4°C. After the incubation, cells were washed thrice with PBS and probed with respective secondary antibodies (Life Technologies) attached with specific fluorophore for 1h at room temperature. For detection of lysosome, 100nM LysoTracker™ Red DND-99 was added in the growing cells for 1h. before fixation. Images were captured using Zeiss LSM800 confocal microscope and analyzed with Imaris7 software. All the interactions between organelles were measured by calculating Pearson’s and Mande’s coefficient of colocalization using coloc plug-in of Imaris7 software.

### Passive avoidance test

First, the rats were subjected to passive avoidance (shuttle box) training. Rodents have a natural tendency to move towards dark. The passive avoidance device had two identical compartments comprising (25 × 25 x 20 cm), which were divided by a guillotine door. One of the chambers are lightened and the other being dark. Electric shocks were conveyed to the grid floor by an isolated stimulator. At the beginning of the experiment rats were habituated in the device for 5 min in the lighted chamber facing away from the entrance to the dark side. On acquisition day the guillotine door was raised, and the latency to enter the dark chamber was recorded. If the animals did not enter the dark compartment within 5 min, they were eliminated from the experiment. Once they entered the dark chamber they received a foot shock of 0.8mA which forced them to return to the light chamber. On the next day defined as the probe day, the same rats were kept in the light chamber and time taken by them to enter the dark chamber was noted as the latency time. If the rats did not enter the dark chamber by 300 s, the successful acquisition of passive avoidance response was said to occur. The experimental work and data recording was performed by a software namely SHUTAVOID by Panlab.

### Novel object recognition (NOR)

The NOR task evaluates the rodents’ ability to recognize a novel object in the environment. The task procedure consists of three phases: habituation, familiarization, and test phase. In the habituation phase, each animal is allowed freely exploring the open-field arena in the absence of any objects. After familiarization phase, a single animal is placed in the open-field arena containing two identical sample objects (A & A), for 5 min. After 24 hr the animal was returned to the open-field arena with two objects, one is identical to the sample and the other one is novel (A & B). Test was performed for 5 min. During both the familiarization and the test phase, objects are located in opposite and symmetrical corners of the arena and location of novel versus familiar object is counterbalanced. The wait latency of the animals observed in each of the objects on probe day was calculated and designated as TN and TF for the novel and familiar objects respectively. Discrimination index (DI) and preference index (PI) were derived according to the formulae:

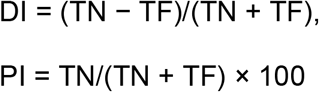

### Elevated Plus maze test

Elevated plus-maze was used for the assessment of memory processes in the animals on 15^th^ day following the method of ^78^ as used in rats. The apparatus was made of a wooden plus-shaped maze elevated 60 cm from the floor having two open arms measuring 50 x 10 cm, crossed at right angles by two arms of the same dimensions enclosed by 50 cm high walls. One cm high plywood edge surrounding the open arms was added to prevent the animals from falling off the maze. A rat was placed on the open arm facing away from the centre and the transfer latency (TL; time in which the rat moves from the open arm to closed arms) was recorded on the day 1 and again on day after. The time taken by the animal, on day 1 to move from open arm to the closed arm is recorded as initial transfer latency (ITL). And that on day 2 as retention transfer latency (RTL).A duration of 5 min was employed for each subject to observe the data.

### Statistical Analysis

Software GraphPad Prism 5.00 (Graph Pad, San Diego) was used to analyze the experiments performed at triplicate unless mentioned otherwise. The P-values were obtained with the help of Non-parametric Student’s t-test. Animal data are represented as means ±SEM and analyzed by one way ANOVA followed by Bonferroni’s test.

Information related to antibodies, primers and miRNA assays are available as Supplemental Table S3-5.

## Acknowledgement

We acknowledge Witold Filipowicz, Gunter Meister and J.M. Backer for different plasmids constructs. SNB is supported by The Swarnajayanti Fellowship from Dept. of Science and Technology, Govt. of India, while D.D., S.G. and I.M. received their support from CSIR, India. We were supported by funds from High Risk High Reward Project Grant, Dept. of Science and Technology, Govt. of India.

## Author contributions

S.N.B. Conceptualize the project, designed research and analyzed data; D.D., I.M., S.G. and R.P. performed research and did animal experiments. S.N.B., D.D. and I.M. along with S.C.B. and S.C. analyzed data; S.N.B., D.D., S.C.B. and I.M.. wrote the paper.

## Appendix Data Expression profile analysis of miRNA and their corresponding target mRNA in cancer

In order to determine whether similar “regulator(s) to target(s)” relationship patterns could be prevalent in other diseases or cell types; we have determined whether these de-regulated mRNA could be related to any other disease condition as well by performing a disease enrichment analysis considering the set of commonly up-regulated mRNA. With the help of disease enrichment analysis, we found that these genes are also associated with cancer (Appendix Figure 1). Therefore, subsequently we have analyzed regulator(s) to target(s)” (miRNA:mRNA) expression profiles in different cancer tissues. A substantial fraction of up-regulated miRNA had their target mRNA as up-regulated in oral squamous cell carcinoma, colorectal and breast cancer as well (Appendix Figure 2A,B,C). In this scenario, we observe that a similar percentage of down-regulated miRNA and up-regulated miRNA have their target mRNAs fractions as up-regulated (Appendix Figure 2D).

---

**Appendix Figure 1:**
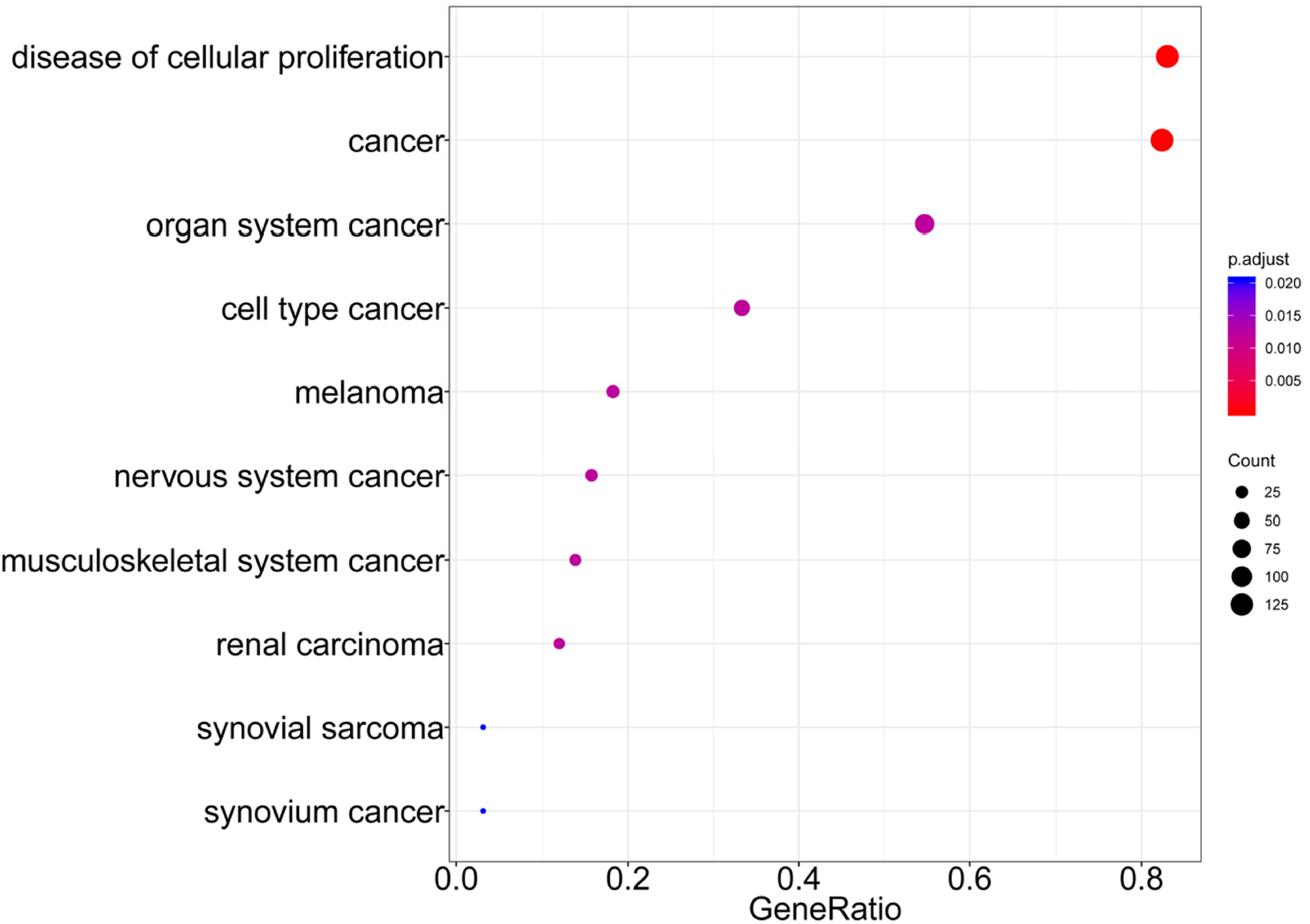
Disease associations of genes up-regulated among multiple brain cortical regions in Alzheimer’s patients. P-value and gene ratio of diseases (top 10) obtained based on enrichment analysis considering the common up-regulated genes among different cortical regions in Alzheimer’s disease patients is shown here.

**Appendix Figure 2:**
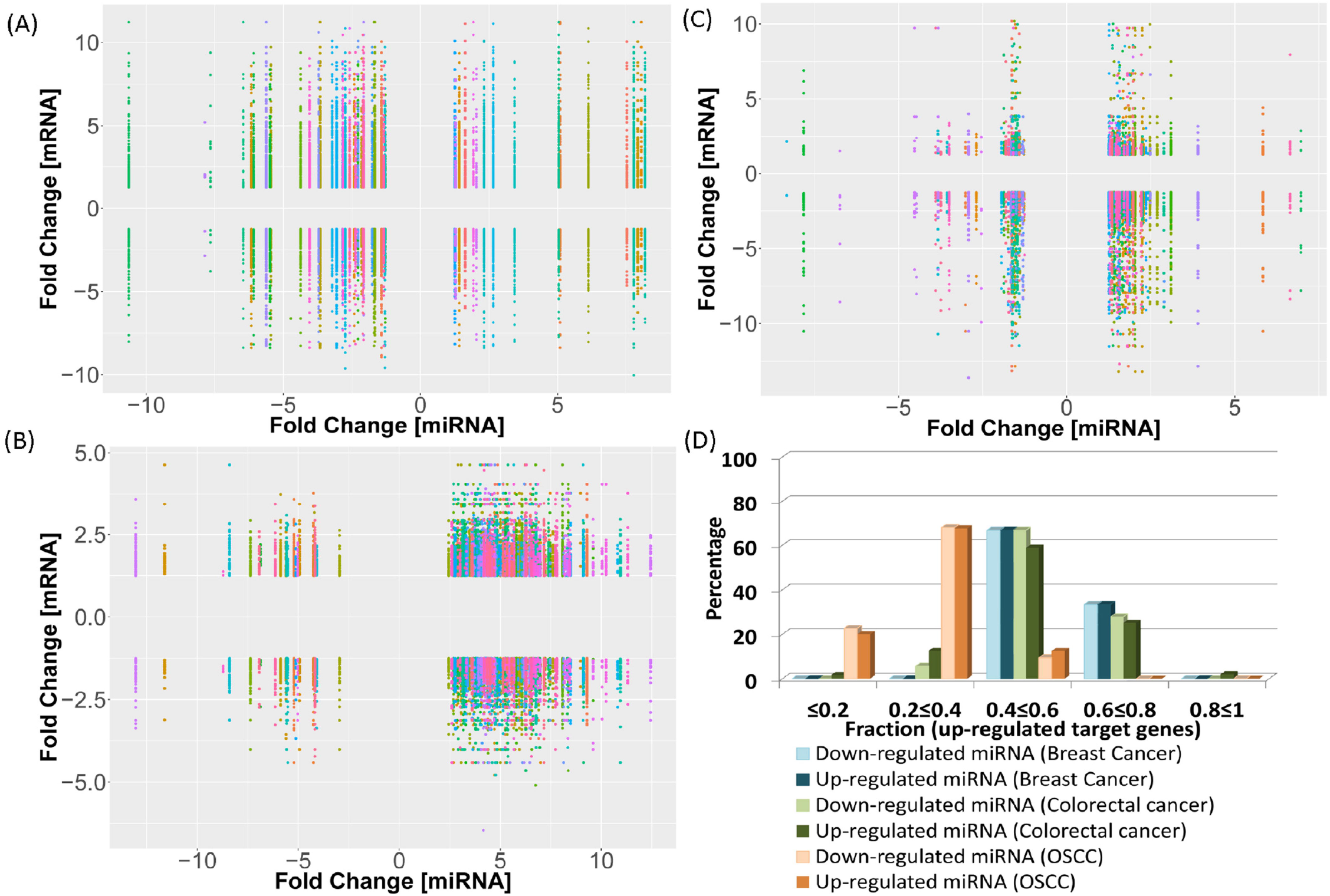
Probable miRNA:mRNA expression relationships in cancer patient samples. (A-C) Fold changes in differentially expressed miRNA and their corresponding differentially expressed target mRNA in oral squamous cell carcinoma (A), colorectal (B) and breast cancer (C) are depicted here. (D) Comparison among percentage of differentially expressed miRNA that have different subsets or fractions of their target mRNA as up-regulated in highly proliferative tissues (different cancer datasets).

