## Supplementary material for "Rheb-mTOR Activation Rescues Amyloid Beta-Induced Cognitive Impairment and Memory Function by Restoring miR-146 Activity in Glial Cells": Sup Figs and Tables

#### Supplementary Figures and Tables

**Supplemental Table S1 Differentially expressed miRNAs in the pre-frontal cortex with their respective fold changes and p-value**

| | miRNA ID | log2FoldChange | P-value |
|---|---|---|---|
| 1 | hsa-miR-100-5p | 1.43 | 1.00E-02 |
| 2 | hsa-miR-141-3p | 1.69 | 9.81E-03 |
| 3 | hsa-miR-142-5p | 1.82 | 1.94E-05 |
| 4 | hsa-miR-146a-5p | 1.29 | 2.25E-02 |
| 5 | hsa-miR-148b-3p | 2.07 | 1.04E-03 |
| 6 | hsa-miR-152 | 1.78 | 4.14E-06 |
| 7 | hsa-miR-153 | 1.67 | 1.99E-07 |
| 8 | hsa-miR-18a-5p | 1.58 | 5.09E-05 |
| 9 | hsa-miR-19a-5p | 1.46 | 1.75E-02 |
| 10 | hsa-miR-208b | 1.65 | 6.30E-08 |
| 11 | hsa-miR-20a-3p | 1.52 | 4.01E-04 |
| 12 | hsa-miR-26a-1-3p | 2.03 | 3.34E-03 |
| 13 | hsa-miR-302a-3p | 2.41 | 3.06E-02 |
| 14 | hsa-miR-302b-3p | 2.35 | 4.78E-03 |
| 15 | hsa-miR-32-5p | 1.27 | 6.25E-07 |
| 16 | hsa-miR-338-3p | 1.32 | 1.84E-06 |
| 17 | hsa-miR-3613-3p | 1.56 | 3.82E-05 |
| 18 | hsa-miR-3676-5p | 1.58 | 4.55E-03 |
| 19 | hsa-miR-374a-5p | 2.15 | 2.18E-04 |
| 20 | hsa-miR-374b-5p | 1.47 | 1.69E-04 |
| 21 | hsa-miR-488-3p | 1.31 | 2.96E-08 |
| 22 | hsa-miR-5000-3p | 1.58 | 1.02E-02 |
| 23 | hsa-miR-500a-5p | 1.44 | 3.68E-02 |
| 24 | hsa-miR-516a-5p | 1.69 | 3.03E-05 |
| 25 | hsa-miR-519a-3p | 1.29 | 1.94E-03 |
| 26 | hsa-miR-590-3p | 1.27 | 2.15E-06 |
| 27 | hsa-miR-675-3p | 2.11 | 4.54E-02 |
| 28 | hsa-miR-7-1-3p | 1.52 | 9.75E-10 |
| 29 | hsa-miR-941 | 2.20 | 3.75E-03 |
| 30 | hsa-miR-99b-5p | 1.46 | 3.69E-03 |
| 31 | hsa-miR-1225-3p | -2.30 | 5.66E-10 |
| 32 | hsa-miR-1225-5p | -1.78 | 1.39E-04 |
| 33 | hsa-miR-1276 | -1.66 | 1.49E-03 |
| 34 | hsa-miR-132-3p | -2.02 | 2.77E-14 |
| 35 | hsa-miR-132-5p | -1.89 | 3.19E-10 |
| 36 | hsa-miR-212-3p | -1.70 | 4.76E-08 |
| 37 | hsa-miR-212-5p | -1.87 | 2.07E-06 |
| 38 | hsa-miR-25-5p | -1.36 | 8.88E-03 |
| 39 | hsa-miR-323a-3p | -1.30 | 3.66E-07 |
| 40 | hsa-miR-323a-5p | -1.53 | 9.33E-06 |
| 41 | hsa-miR-3607-3p | -2.76 | 2.69E-10 |
| 42 | hsa-miR-3651 | -1.95 | 4.09E-02 |
| 43 | hsa-miR-3653 | -2.08 | 4.96E-07 |
| 44 | hsa-miR-376a-5p | -1.49 | 1.97E-05 |

| | | | |
|---|---|---|---|
| 45 | hsa-miR-4284 | -1.43 | 2.11E-02 |
| 46 | hsa-miR-4520a-3p | -1.71 | 3.28E-03 |
| 47 | hsa-miR-505-5p | -1.43 | 1.92E-02 |
| 48 | hsa-miR-511 | -1.85 | 1.51E-02 |
| 49 | hsa-miR-5701 | -2.23 | 1.97E-08 |
| 50 | hsa-miR-6087 | -2.56 | 1.54E-03 |
| 51 | hsa-miR-6721-5p | -2.39 | 9.94E-03 |
| 52 | hsa-miR-877-5p | -1.52 | 1.94E-07 |
| 53 | hsa-miR-885-3p | -2.15 | 9.54E-15 |

**Supplemental Table S2 Expression profile of miR-146a-5p target genes in brain cortical regions of Alzheimer's disease patients.** Fold change and p-values of differentially expressed miR-146a-5p within cortical regions of Alzheimer's disease patients determined based on differential expression analysis of mRNA profiling data is represented here.

| Frontal cortex | | | Superior frontal gyrus | | | Dorsolateral pre-frontal cortex | | |
|---|---|---|---|---|---|---|---|---|
| **Target gene name** | **Fold change** | **p-value** | **Target gene name** | **Fold change** | **p-value** | **Target gene name** | **Fold change** | **p-value** |
| brca2 | 1.27 | 8.96E-03 | cxcr4 | 4.45 | 5.39E-05 | tlr4 | -2.34 | 5.96E-03 |
| cdkn1a | 1.27 | 3.93E-09 | egfr | 1.89 | 5.03E-03 | wasf2 | -1.44 | 2.92E-02 |
| egfr | 1.48 | 1.36E-03 | erbb4 | 2.76 | 4.37E-07 | notch2 | -1.58 | 2.12E-02 |
| tlr4 | 1.27 | 3.03E-08 | tlr4 | 1.33 | 2.16E-03 | lfng | -1.75 | 2.13E-02 |
| slpi | 1.64 | 1.71E-03 | stat1 | -1.42 | 2.39E-02 | tgfb1 | -2.57 | 2.32E-02 |
| card10 | 1.45 | 6.02E-11 | card10 | 1.39 | 4.72E-02 | klf4 | -3.34 | 2.67E-03 |
| icam1 | 1.40 | 2.87E-02 | cops8 | -1.26 | 4.64E-02 | tcf7 | -1.92 | 1.15E-02 |
| ccl5 | 1.62 | 5.28E-06 | elavl1 | 2.11 | 3.86E-03 | col18a1 | -2.04 | 3.82E-02 |
| cnot6l | 1.62 | 1.25E-02 | casp7 | 1.69 | 3.86E-03 | prkacb | 1.66 | 4.21E-02 |
| cpm | 1.71 | 1.59E-13 | notch2 | 2.26 | 1.48E-03 | igf1 | 2.91 | 4.39E-03 |
| tgfb1 | 1.72 | 5.13E-03 | sox2 | 2.11 | 9.91E-04 | slc3a1 | 2.20 | 2.66E-02 |
| fancm | 2.94 | 2.61E-02 | lfng | 1.74 | 2.26E-03 | cpt1a | -1.32 | 3.16E-02 |
| notch1 | 1.34 | 1.29E-19 | mif | -1.81 | 4.21E-02 | lrp5 | -2.88 | 1.22E-02 |
| nos2 | 1.49 | 4.48E-03 | notch1 | 1.33 | 1.20E-03 | rassf3 | -1.67 | 3.20E-02 |
| egr1 | -2.05 | 9.95E-13 | nfat5 | 1.55 | 3.48E-02 | bcat2 | -1.94 | 2.45E-02 |
| tgif1 | 1.40 | 5.36E-04 | med1 | 1.39 | 1.10E-02 | necab1 | 2.11 | 1.77E-02 |
| snap25 | -3.55 | 7.86E-26 | egr1 | -1.95 | 7.87E-03 | znrf3 | -1.32 | 1.50E-02 |
| st6gal2 | 1.46 | 1.39E-02 | camk2d | -1.47 | 9.12E-03 | maff | -2.54 | 3.95E-02 |
| tmprss5 | 1.78 | 4.29E-11 | snap25 | -4.50 | 1.35E-04 | gpr146 | -1.88 | 4.60E-02 |
| shcbp1 | 1.28 | 6.20E-11 | klf4 | 2.49 | 1.28E-02 | gpr116 | -1.57 | 5.62E-03 |
| rhobtb3 | 1.42 | 2.96E-16 | mettl7a | 1.31 | 3.75E-04 | eid1 | 1.75 | 1.63E-02 |
| mical2 | -1.62 | 9.70E-24 | kctd15 | 1.85 | 1.54E-02 | pigb | 1.41 | 3.53E-02 |
| plekhg5 | 2.38 | 2.25E-02 | rhobtb3 | 2.34 | 9.71E-03 | tpd52 | 1.56 | 1.38E-02 |
| lbr | 1.50 | 1.31E-03 | ppp1r11 | -1.46 | 3.47E-03 | sox4 | -1.63 | 3.19E-02 |
| cd93 | 1.43 | 8.11E-09 | mical2 | -1.52 | 4.29E-02 | rhpn2 | -2.20 | 2.83E-03 |
| sgk3 | 1.53 | 5.53E-12 | copa | -1.33 | 4.03E-02 | btbd3 | 1.29 | 3.77E-02 |
| hormad2 | 2.32 | 1.20E-03 | pacs2 | 1.28 | 1.40E-03 | pla1a | -4.08 | 3.65E-03 |
| alg10b | 1.90 | 4.30E-03 | cybrd1 | 1.78 | 1.31E-02 | fli1 | -1.86 | 3.39E-02 |
| tcf7 | 1.45 | 2.54E-05 | brwd1 | 1.54 | 7.17E-05 | synj1 | 1.58 | 4.35E-02 |
| mid1ip1 | 1.37 | 3.61E-15 | kat2b | 1.68 | 7.78E-03 | midn | -1.89 | 2.07E-02 |
| kif11 | -3.94 | 2.75E-02 | Sep-07 | -2.00 | 2.73E-03 | znf768 | -1.61 | 2.73E-02 |
| cd274 | 1.88 | 4.86E-04 | braf | -1.33 | 3.25E-02 | metrnl | -2.40 | 3.26E-02 |
| hnf4a | 1.38 | 2.52E-02 | cse1l | -2.28 | 4.27E-03 | aen | -2.01 | 4.39E-03 |
| ypel2 | 1.43 | 8.66E-03 | crot | -1.30 | 1.05E-02 | stk40 | -1.80 | 1.09E-02 |
| plk2 | -2.95 | 5.39E-21 | insig2 | -1.29 | 4.06E-02 | itpripl2 | -1.99 | 2.38E-02 |
| il6st | 1.38 | 1.42E-02 | hlf | -1.72 | 1.38E-02 | pcdhgb6 | -1.89 | 3.11E-02 |
| c6 | 2.40 | 1.54E-03 | smurf2 | 1.31 | 1.97E-04 | | | |
| zbtb20 | 1.48 | 2.66E-20 | ube2b | -1.62 | 1.93E-02 | | | |
| cpt1a | 1.46 | 8.40E-04 | xbp1 | 2.01 | 3.44E-04 | | | |
| lrp5 | 1.40 | 5.50E-14 | plk2 | -2.49 | 1.42E-02 | | | |

| | | | | | |
|---|---|---|---|---|---|
| ablim1 | 1.31 | 1.02E-02 | serinc5 | 1.29 | 4.75E-03 |
| mettl7b | 1.57 | 4.35E-13 | arf4 | -1.58 | 3.46E-05 |
| creb1 | 1.32 | 4.52E-11 | samm50 | -1.65 | 1.01E-03 |
| cdc73 | -1.38 | 6.84E-03 | zbtb20 | 2.92 | 2.29E-04 |
| fam107b | 1.38 | 2.65E-12 | usp25 | -1.40 | 4.23E-02 |
| hsd17b7 | 1.34 | 4.82E-10 | slc3a1 | -1.46 | 3.17E-02 |
| atrn | -1.36 | 9.68E-17 | ablim1 | 1.56 | 9.72E-03 |
| rbl1 | 1.41 | 9.21E-05 | atp5b | -2.15 | 3.13E-02 |
| clta | -2.97 | 6.19E-17 | tm9sf2 | -1.84 | 8.74E-03 |
| slc6a13 | 1.96 | 1.66E-13 | mrpl30 | -1.50 | 1.96E-03 |
| psma1 | -1.39 | 1.08E-02 | glul | 2.14 | 1.77E-02 |
| tirap | 1.53 | 1.17E-03 | fam107b | 1.46 | 3.43E-02 |
| ddx6 | 2.49 | 6.30E-05 | camsap1 | -1.41 | 2.77E-02 |
| pank1 | 1.52 | 3.67E-03 | cers6 | -2.30 | 8.53E-05 |
| atp6v1h | -1.56 | 1.79E-23 | sp3 | 2.05 | 4.70E-04 |
| hsd17b13 | 1.63 | 1.08E-07 | ndrg3 | -1.57 | 6.69E-03 |
| ror1 | 1.30 | 6.14E-14 | ralgapb | -1.29 | 1.84E-02 |
| ago2 | 1.70 | 1.91E-14 | dhx36 | -1.50 | 1.63E-02 |
| rcor1 | 2.01 | 4.13E-02 | pnisr | 1.74 | 3.90E-02 |
| necab1 | -2.80 | 2.37E-23 | clta | -1.44 | 3.60E-02 |
| maff | 2.20 | 8.69E-04 | akr1a1 | -1.54 | 7.02E-03 |
| serpinb9 | 1.91 | 1.28E-04 | psma1 | -2.40 | 6.68E-04 |
| tnrc6a | 1.33 | 8.46E-08 | nek1 | 1.97 | 1.47E-03 |
| ddx17 | 1.79 | 7.05E-08 | atp6v1h | -2.39 | 2.59E-02 |
| cfhr1 | 1.31 | 7.76E-03 | dhcr24 | -2.39 | 3.87E-04 |
| rbm47 | 1.46 | 9.45E-07 | srrm2 | 2.02 | 4.06E-02 |
| ccr5 | 1.28 | 2.98E-07 | necab1 | -1.67 | 2.51E-02 |
| slc5a3 | 2.54 | 1.15E-09 | znrf3 | 2.15 | 3.77E-04 |
| ndc1 | 1.29 | 7.13E-08 | maff | 1.47 | 1.72E-03 |
| cd84 | 1.26 | 1.52E-03 | maml1 | 1.52 | 5.06E-03 |
| rora | 1.40 | 9.37E-05 | hnrnpc | -1.39 | 6.10E-03 |
| stxbp2 | 1.36 | 4.81E-09 | dlgap4 | 1.50 | 7.99E-03 |
| itch | 1.39 | 6.78E-09 | rab18 | -1.59 | 1.90E-03 |
| znf264 | 1.36 | 1.97E-12 | slc5a3 | 1.56 | 1.79E-02 |
| ddhd1 | -1.86 | 4.45E-03 | eid1 | -1.91 | 1.41E-02 |
| hm13 | 1.58 | 9.86E-05 | rev3l | 1.47 | 2.22E-02 |
| akt2 | 3.32 | 1.28E-03 | gopc | -1.86 | 2.22E-03 |
| slc1a5 | 1.50 | 9.02E-14 | kmt2c | 1.29 | 2.54E-03 |
| fbxo3 | -1.67 | 2.07E-10 | rora | 1.53 | 3.23E-02 |
| dnph1 | 2.00 | 1.25E-04 | tpd52 | -2.47 | 1.36E-03 |
| srsf11 | 1.48 | 1.04E-08 | gsk3b | -1.32 | 2.44E-02 |
| mkln1 | 1.30 | 8.40E-19 | papola | 1.27 | 3.14E-02 |
| gtpbp3 | 1.43 | 1.45E-04 | supt16h | -1.28 | 1.18E-03 |
| btbd3 | 1.32 | 6.22E-03 | cecr2 | 2.34 | 5.76E-04 |
| ccnb1 | -1.67 | 9.49E-14 | chmp4b | -1.39 | 3.30E-02 |
| gltp | 2.12 | 1.29E-18 | mid1 | 1.65 | 1.16E-02 |

| | | | | | |
|---|---|---|---|---|---|
| pla1a | 1.63 | 9.92E-04 | akt2 | 1.35 | 1.02E-02 |
| tnfaip8 | 1.81 | 4.37E-10 | card8 | 1.64 | 4.37E-04 |
| nacc2 | 1.44 | 3.42E-02 | grpel1 | -1.66 | 7.79E-03 |
| hnrnpu | 1.31 | 1.47E-04 | fbxo3 | -1.44 | 1.51E-02 |
| slc26a2 | 2.69 | 3.00E-03 | slc38a1 | -1.39 | 3.76E-03 |
| elk4 | 1.44 | 3.37E-02 | aak1 | -2.10 | 9.46E-03 |
| synj1 | 2.16 | 7.42E-04 | pds5a | -1.34 | 1.14E-02 |
| sqstm1 | 1.68 | 3.98E-16 | mkln1 | 1.36 | 1.48E-02 |
| hipk1 | 1.80 | 5.20E-06 | arpp19 | -2.82 | 5.90E-05 |
| slc4a1ap | -1.60 | 9.21E-15 | phf20l1 | -1.28 | 2.74E-03 |
| tp53inp1 | 1.35 | 3.14E-18 | clip1 | 1.56 | 1.94E-02 |
| znf367 | 1.73 | 3.19E-03 | rhpn2 | 2.45 | 5.04E-05 |
| midn | 1.55 | 5.59E-15 | anxa7 | -1.68 | 3.39E-03 |
| zdhhc16 | 1.38 | 5.82E-03 | gltp | 1.29 | 2.57E-02 |
| klhl15 | 1.46 | 2.18E-04 | rfx5 | -1.51 | 2.62E-02 |
| zbtb33 | 1.64 | 1.38E-03 | abracl | -2.02 | 2.17E-03 |
| znf620 | 1.56 | 2.98E-06 | nacc2 | 2.64 | 3.87E-03 |
| efna5 | 2.00 | 2.52E-02 | eif4ebp2 | 1.44 | 2.45E-04 |
| baz1a | 1.69 | 2.63E-06 | aldoa | -1.26 | 3.53E-02 |
| ltb4r2 | 1.36 | 9.44E-04 | nek7 | 1.48 | 1.78E-02 |
| txnip | 1.78 | 1.79E-15 | fli1 | 1.33 | 1.17E-03 |
| rassf5 | 1.72 | 2.93E-05 | uhmk1 | -2.06 | 3.05E-03 |
| | | | carhsp1 | 2.17 | 2.57E-03 |
| | | | azin1 | -1.54 | 4.60E-02 |
| | | | rer1 | -1.95 | 4.14E-06 |
| | | | synj1 | -1.60 | 4.91E-02 |
| | | | sqstm1 | 2.22 | 8.84E-04 |
| | | | slc4a1ap | -1.47 | 8.42E-03 |
| | | | tp53inp1 | 2.36 | 9.87E-04 |
| | | | kbtbd6 | -1.30 | 3.83E-02 |
| | | | usp54 | 1.53 | 4.09E-03 |
| | | | tmem41b | -1.45 | 3.51E-03 |
| | | | znf597 | -1.44 | 3.56E-03 |
| | | | clic4 | 1.34 | 2.14E-02 |
| | | | jun | 1.58 | 4.47E-02 |
| | | | ptar1 | 1.53 | 1.07E-02 |
| | | | peli1 | 1.30 | 4.59E-02 |
| | | | ythdf2 | -1.45 | 2.15E-02 |
| | | | baz1a | 1.73 | 4.95E-04 |
| | | | itpripl2 | 2.21 | 3.60E-03 |
| | | | sft2d2 | 2.07 | 1.26E-03 |

**Supplemental Table S3 Details of Antibodies used for western blot (WB), Imunohistochemistry (IHC) and Immunocytochemistry (ICC)**

| Antibody Name | Raised In | Dilution | Source |
|---|---|---|---|
| GFAP | Mouse | IHC/ICC-1:50 | Sigma Aldrich |
| $A\beta_{1-42}$ | Rabbit | IHC-1:100, ICC/IHC-1:100 | Abcam |
| Ago2 (eIF2C2) | Mouse | WB-1:1000 | Abnova |
| HA | Rat | WB-1:1000 | Roche |
| Dcp1 | Mouse | WB-1:1000, ICC/IHC-1:100 | Novus |
| Rck/p54 | Rabbit | WB-1:10000, ICC/IHC-1:1000 | Bethyl |
| Alix | Mouse | WB-1:200 | Santa cruz |
| Calnexin | Rabbit | WB-1:10000 | Bethyl |
| HRS | Rabbit | WB-1:1000 | Bethyl |
| β-Actin | Mouse Monoclonal (HRP Conjugated) | WB-1:10000 | Sigma Aldrich |
| Rab7 | Rabbit | WB-1:1000, ICC-1:100 | Cell Signalling |
| DICER | Rabbit | WB-1:5000 | Bethyl |
| Rab5 | Rabbit | WB-1:1000 | Cell Signalling |
| RILP | Goat | WB-1:250 | Santa cruz |
| p-mTOR | Rabbit | WB-1:1000 | Cell Signalling |
| p70-S6K | Rabbit | WB-1:1000 | Cell Signalling |
| mTOR | Rabbit | WB-1:1000, ICC-1:100 | Cell Signalling |
| 4G-10 | Mouse | WB-1:1000 | Millipore |
| Myc | Mouse | WB-1:1000 | Cell Signalling |

**Supplemental Table S4 mRNA Primers used for quantification**

| Target | 5'Forward Primer3' | 5'Reverse Primer3' |
|---|---|---|
| RL | CCAAGCAAGATCATGC | GCTCTTGATGTACTTACCC |
| Pre-miR-122 | CCTTAGCAGAGCTGTGGAG | GCCTAGCAGTAGCTATTTAG |
| 18SrRNA | TGACTCTAGATAACCTCGGG | GACTCATTCCAATTACAGGG |
| IL-1β | GTGGATCCCAAACAATACCC | AACTATGTCCCGACCATTGC |
| IL-6 | TACCCCAACTTCCAATGCTC | ACCACAGTGAGGAATGTCCA |
| GAPDH | CAGGGGGGAGCCAAAAGGG | CTTGGCCAGGGGTGCTAAGC |

**Supplemental TableS5 Details of miRNA primers used for Taqman based quantification**

| miRNA Name | ASSAY ID |
|---|---|
| Let-7a | 000377 |
| miR-9 | 001089 |
| miR-16 | 000391 |
| miR-21 | 000397 |
| miR-24 | 000402 |
| miR-29a | 000412 |
| miR-33-5p | 002135 |
| miR-101 | 000438 |
| miR-122 | 000445 |
| miR-125b | 000449 |
| miR-128a | 002216 |
| miR-142-3p | 000464 |
| miR-145 | 002278 |
| miR-146a | 000468 |
| miR-155 | 002571 |
| miR-181c | 000482 |
| miR-184 | 000485 |
| U6 SnRNA | 001973 |

### Supplementary Figure S1

**A**

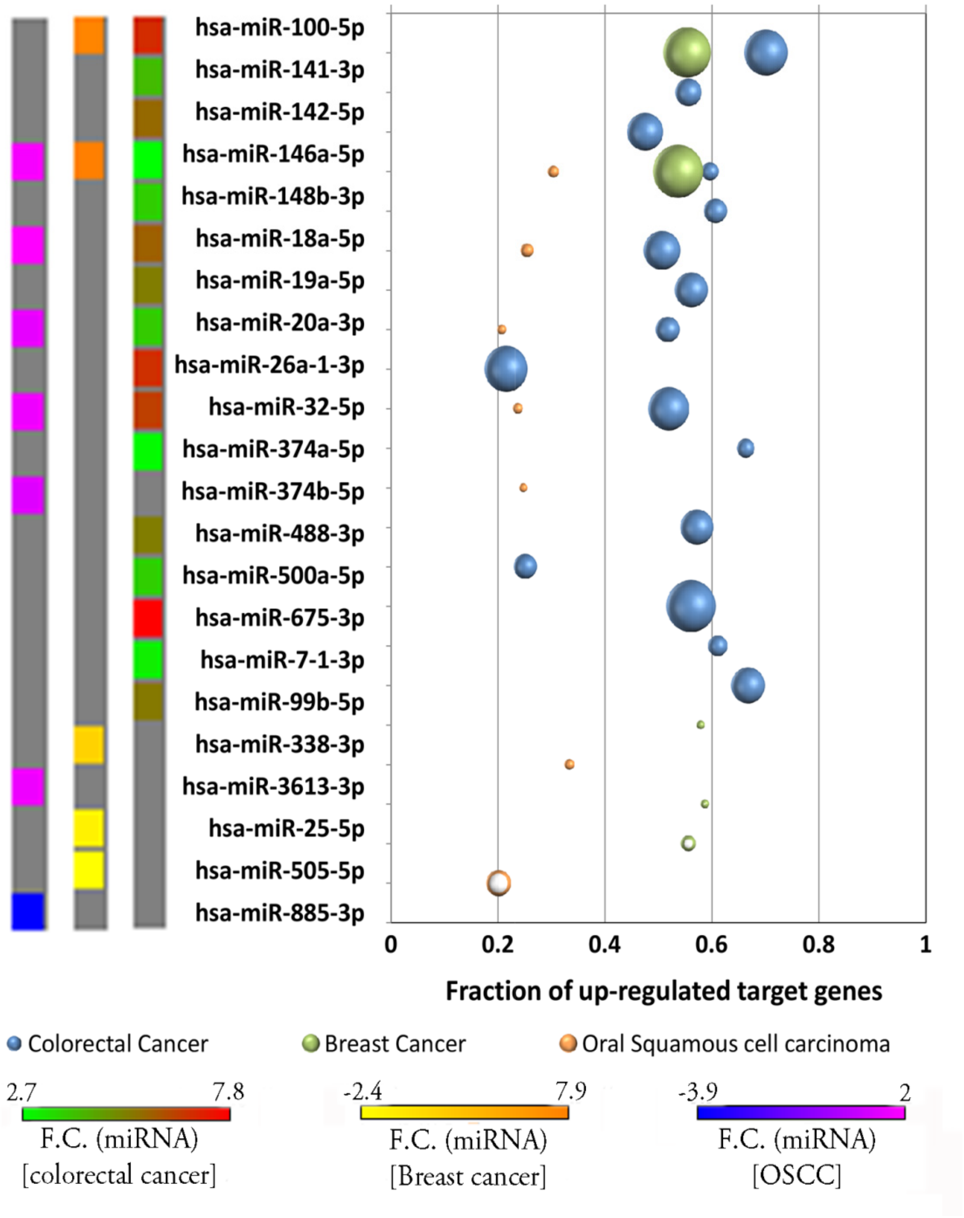

**Figure S1 Probable regulator(s) to target(s) [miRNA:mRNA] expression relationships in Cancer.**

The possible relationship between miRNA and their target mRNA in different cancer tissues was studied considering commonly differentially expressed miRNA in AD and cancer. The fraction of up-regulated target mRNA of these differentially expressed miRNA in colorectal cancer, breast cancer and oral squamous cell carcinoma is shown here.

Supplementary Figure S2

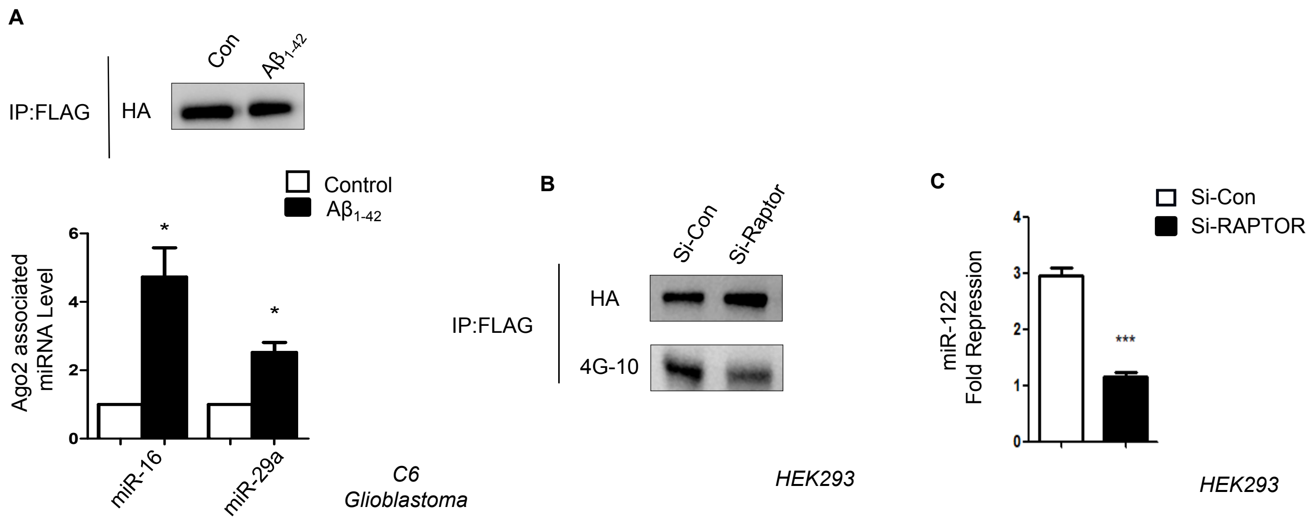

**Figure S2 RAPTOR knockdown decreases miRNA activity and Ago2 phosphorylation**

**A** Graphs showing level of Ago2 associated miR-16 and miR-29a level upon $A\beta_{1-42}$ treatment in C6 glioblastoma cells. qPCR data was normalized with the amount of Ago2 pulled down from IP reaction.

**B** Western blot data showing amount of Ago2 phosphorylation and the amount of Ago2 pulled down from Si-con and Si-RAPTOR transfected HEK293 cells stably expressing F-HA-Ago2.

**C** Effect of RAPTOR knockdown on miR-122 activity in HEK293 cells. Dual Luciferase assay showing fold repression of exogenously expressed miR-122 in both control and RAPTOR knockdown cell.

For statistical significance, minimum three independent experiments were considered in each case unless otherwise mentioned and error bars are represented as mean ± S.E.M. P-values were calculated by utilizing Student's t-test. ns: non-significant, *P < 0.05, ***P < 0.0001.

### Supplementary Figure S3

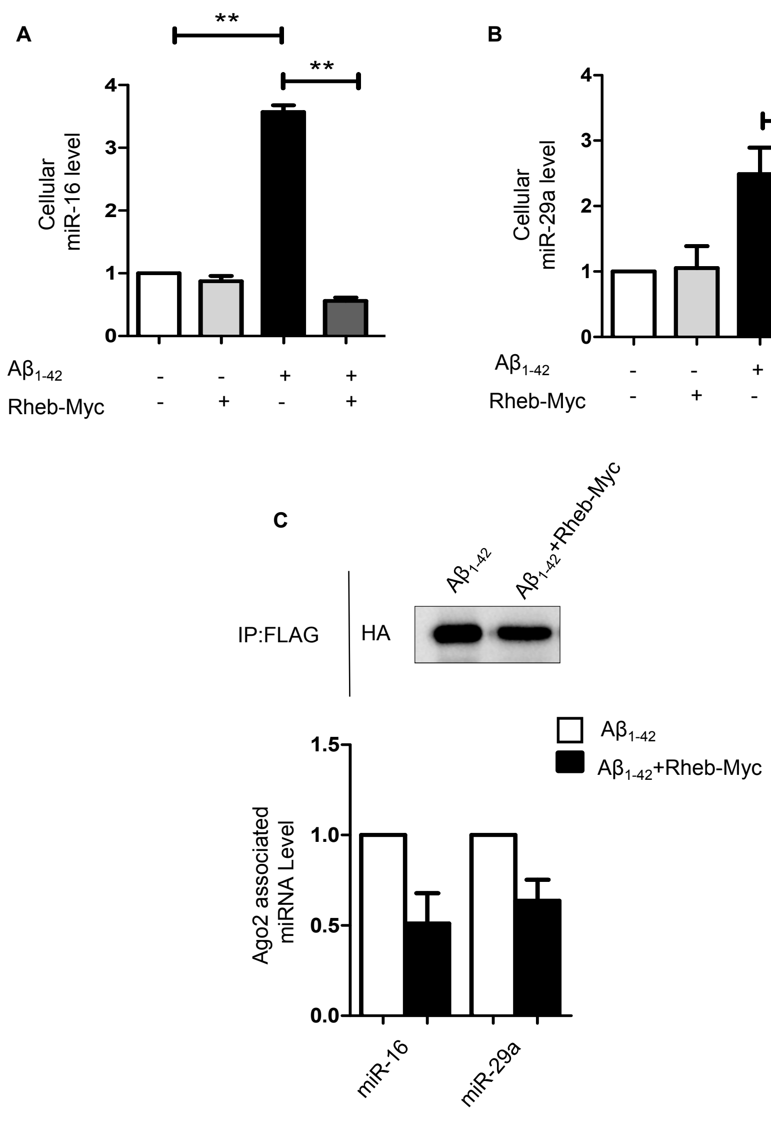

**Figure S3 Rheb-myc expression decreases miRNP formation in A$\beta_{1\text{-}42}$ treated cells**

**A-B** Graphs showing levels of miR-16 and miR-29a miRNA in both control vector and Rheb-myc expressing cells exposed with A$\beta_{1\text{-}42}$. qPCR data was normalized with U6 snRNA.

**C** Graphs depicting Ago2 associated miR-16 and miR-29a level in both control vector and Rheb-myc expressing cells exposed with A$\beta_{1\text{-}42.}$ qPCR data was normalize with the amount of Ago2 pulled down from IP reaction.

For statistical significance, minimum three independent experiments were considered in each case unless otherwise mentioned and error bars are represented as mean ± S.E.M. P-values were calculated by utilizing Student's t-test. ns: non-significant, *P < 0.05, **P < 0.01.

### Supplementary Figure S4

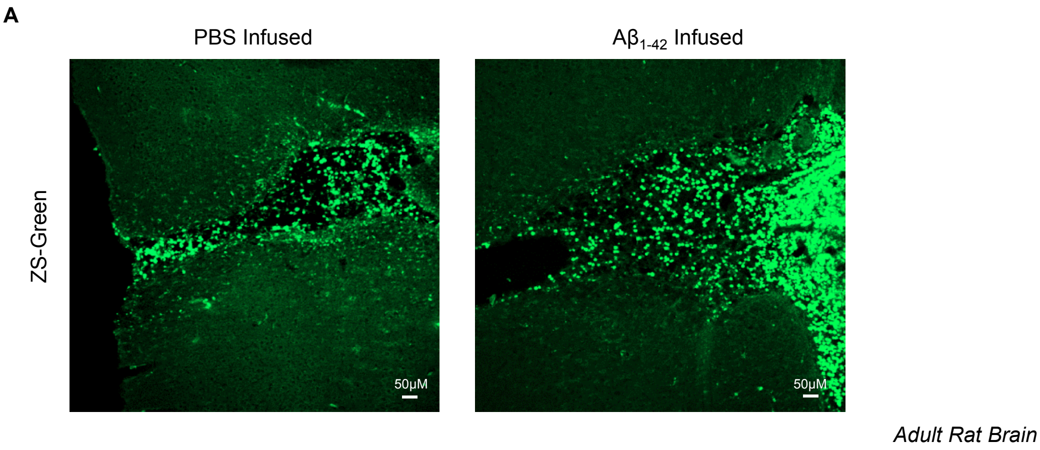

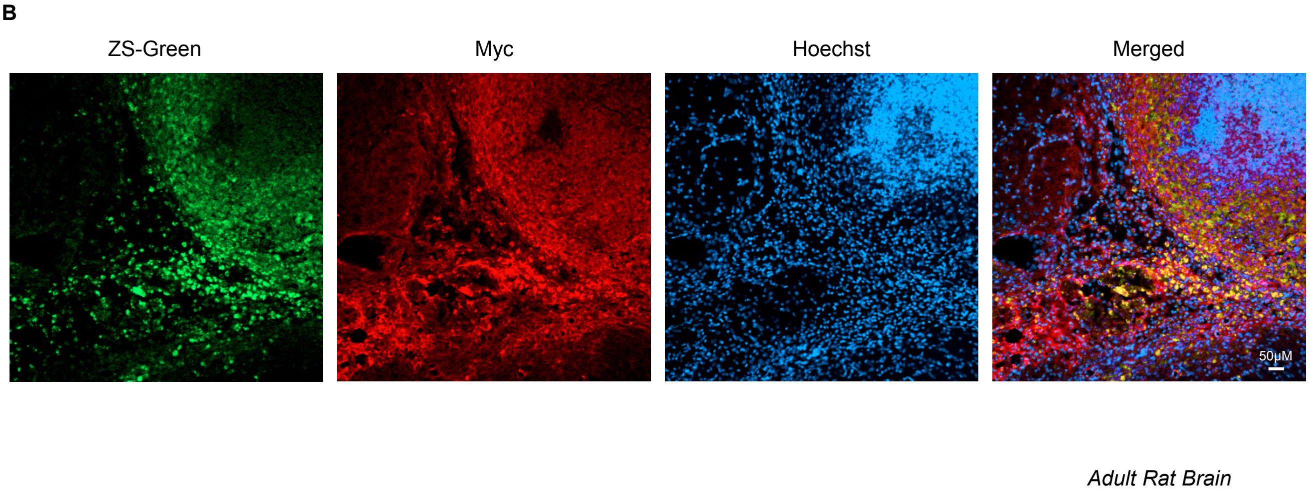

**Figure S4 Confirmation of Rheb-Myc expression in adult rat brain. The protein expression plasmid was co-infused along with $A\beta_{1-42}$ oligomer in adult rat brain.**

**A** Immunohistochemistry panels are showing brain sections that were infused either with PBS or $A\beta_{1-42}$ oligo along with ZS-Green expression vector.

**B** Confocal images showing immunohistochemistry sections with ZS-Greenand Rheb-Myc expression vector. The green color positive cells that are depicting ZS-Green expression were also stained for Rheb-Myc and dual coloured cells were visualized as yellow.
